# Accounting for stimulations that do not elicit motor-evoked potentials when mapping cortical representations of multiple muscles

**DOI:** 10.1101/2021.07.29.454279

**Authors:** Fang Jin, Sjoerd M Bruijn, Andreas Daffertshofer

## Abstract

The representation of muscles in the cortex can be mapped using navigated transcranial magnetic stimulation. The mapping can be quantified via measures like the centre of gravity or the size of the region of excitability. Determining these measures typically relies only on stimulation points that yield motor-evoked potentials (MEPs); stimulations that do not elicit an MEP, i.e., non-MEP points, are ignored entirely. In this study, we show how incorporating non-MEP points may affect the estimates of the size and centroid of the excitable area in eight hand and forearm muscles after mono-phasic single-pulse TMS. We performed test-retest assessments in twenty participants and estimated the reliability of centroids and sizes of the corresponding areas using inter-class correlation coefficients. For most muscles, the reliability turned out good, if not excellent. As expected, removing the non-MEP points significantly decreased area sizes and area weights, suggesting that conventional approaches that do not account for non-MEP points are likely to overestimate the regions of excitability.

## Introduction

Single-pulse transcranial magnetic stimulation (TMS) is a non-invasive and painless technique that monitors neurophysiological alterations of the human motor cortex (Barker et al., 1985; Schambra et al., 2003). A TMS coil discharge at suitable intensity will induce transient currents and cause depolarisation of axons of nerve cells (Rossini et al., 2015). When applied over the motor cortex, this can elicit a motor-evoked potential (MEP) that can be recorded in contralateral target muscles using conventional electromyography (EMG). Amplitudes and latencies of the MEPs reveal the excitability and conduction times of the cortical-spinal tract. Both have been conceived as valid outcomes of TMS motor mapping (Rossini et al., 1994). Neuroscientists and physicians alike utilised TMS motor mapping to evaluate muscle synergies and motor cortical plasticity (Siebner and Rothwell, 2003), to plan brain tumour surgery (Krieg et al., 2012), or to follow recovery after stroke (Mark et al., 2006; Sondergaard et al., 2021). There is ample evidence that the location at which TMS elicits the maximum MEP is particularly close to the location found using direct cortical stimulation, which is considered the gold standard in motor mapping. The localisation clearly outperforms other modalities like magnetoencephalography (Tarapore et al., 2012) or functional magnetic resonance imaging (Forster et al., 2011).

TMS combined with neuro-navigation increases mapping accuracy (Krieg et al., 2017). A popular approach for coil positioning is a pseudo-random walk. Delivering stimulations at random locations roughly evenly spaced over the motor cortex is more efficient and potentially more accurate than time-consuming, course-grained grid-based positioning. Likely, this random placement will also elicit MEPs in muscles other than the target muscle. Often considered a confounder, this is – in fact – particularly useful when multiple muscles are being evaluated, presuming that the muscles have similar resting motor thresholds (Krieg et al., 2017) and close-by cortical representations (Schieber, 2001). Very recently, Tardelli et al. (2021) used the pseudo-random walk method to assess the cortical representation of abductor digiti minimi, flexor carpi radialis and flexor pollicis brevis. As a rule of thumb: the more muscles are measured simultaneously, the more efficient assessments via pseudorandom coil positioning can be.

Irrespective of the experimental protocol, navigated TMS derived cortical map outcomes should have good reliability (Novikov et al., 2018). Nonetheless, “even most commonly used outcomes such as areas, volumes, the location of centres of gravity (CoGs), and hotspots have [hardly] been validated for being reliable measures in test-retest studies (Kraus and Gharabaghi, 2016)”. We slightly modified this quote from (Novikov et al., 2018), because they and other likewise recent reports did indeed test for the reliability of navigated TMS outcomes considered in the respective studies. For instance, Nazarova et al. (2021) evaluated the reliability of the CoG and the size of the area (volume) of excitability, next to the position of the MEP hotspot. In a grid-based approach, all measures displayed high relative but low absolute reliability, with the latter arguably reflecting between-subject variability.

The area of excitability can be defined as the cortical region within which TMS elicits an MEP. It is usually determined by projecting the focal point of the coil’s magnetic field on a (re-)constructed spherical surface or volume and determining the resulting convex hull. State-of-the-art fine-tuning of this approach is to a priori concentrate on the cortical patch of interest. However, a mapping that agrees with the ‘real’ anatomical structure generally provides better area estimates. This is particularly true when realising that the grey matter border may have large curvatures along gyral ridges (Van Essen, 2004), where spherical approximations will be poor. We followed these lines and extracted subject-specific cortical surfaces at high resolution, projected the stimulation points to that surface and estimated the area spanned by the pair-wise shortest paths connecting the stimulation points. More importantly, we also projected the stimulation points where TMS did *not* elicit an MEP and removed these points from the estimated area. As will be shown, together these steps circumvent potential over-estimation of the area of excitability.

## Methods

### Participants

Twenty healthy, right-handed volunteers (average age: 29.6 ± 7.5, eight females) participated in the study. Prior to the experiment, all participants were screened for contraindications of MRI and TMS through questionnaires (Rossi et al., 2011). All of them provided signed informed consent prior to joining the experimental sessions. The Edinburgh Handedness Inventory served to determine hand dominance (Oldfield, 1971). The study had been approved by the medical ethics committee of Amsterdam University Medical Center (VUmc, 018.213 - NL65023.029.18).

### Materials

Our set-up consisted of three devices: a TMS system, an EMG amplifier, and a neural navigation system. Single-pulse TMS was delivered by a Magstim 200^2^ stimulator (Magstim Company Ltd., Whitland, Dyfed, UK) using a figure-of-eight coil with 70 mm windings. Eight bipolar EMG signals were recorded using a 16-channel EMG amplifier (Porti, TMSi, Oldenzaal, the Netherlands) and continuously sampled at a rate of 2 kHz. The EMG recordings were triggered by the TMS to allow for online EMG assessments using a custom-made Labview-programme with embedded Matlab functions (designed at our department using Labview 2016, National Instruments, Austin, TX, and Matlab 2018b, The MathWorks, Natick, MA). In brief, upon receiving a trigger, peak-to-peak amplitudes and latencies of MEPs were estimated from all EMG signals during the following 500 ms. These outcomes, as well as the original EMG signals (duration = 500 ms), were sent to the neural navigation system (Neural Navigator, Brain Science Tools, De Bilt, The Netherlands, www.brainsciencetools.com) for online monitoring and storage. The neural navigation software also stored the position and orientation of the coil with respect to the head.

Prior to running the TMS protocol, we acquired the participants’ anatomical T1-weighted MRI (3 Tesla Philips Achieva System, Philips, Best, The Netherlands; matrix size 256 × 256 × 211, voxel size 1.0 × 1.0 × 1.0 mm^3^, TR/TE 6.40/2.94 ms). For online neuro-navigation, grey matter was segmented using SPM (https://www.fil.ion.ucl.ac.uk/spm/software/spm12/); note that for offline analysis, we employed a more detailed segmentation via Freesurfer (https://surfer.nmr.mgh.harvard.edu/fswiki, version 7) – see below.

We considered the first dorsal interosseous (FDI), abductor digiti minimi (ADM), abductor pollicis brevis (APB), flexor pollicis brevis (FPB), extensor digitorum communis (EDC), flexor digitorum superficialis (FDS), extensor carpi radialis (ECR) and flexor carpi radialis (FCR) muscles, which were measured using bipolar electrodes (Blue Sensor N-00-S, Ambu, Ballerup, Denmark), placed after cleaning the skin with alcohol; cf. Figure 1. The ground electrode was attached to the ulnar styloid process. We monitored and kept the electrode impedance below 5 kΩ. During the experiment, the orientation of the TMS coil was held 45 degrees to the sagittal plane, and tangential to the scalp. By this, we meant to induce currents in the cortex along the posterior-to-anterior direction. To control the TMS output, we used the Matlab-toolbox Rapid2 (https://github.com/armanabraham/Rapid2).

**Figure 1.**
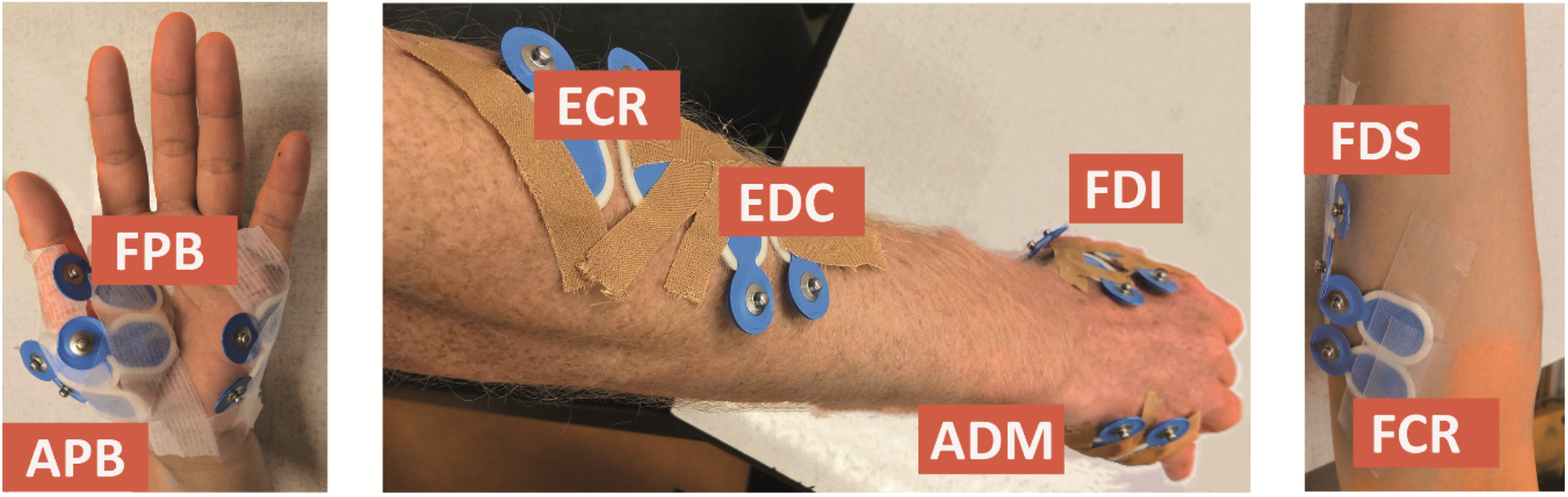
Electrode placement for the first dorsal interosseous (FDI), abductor digiti minimi (ADM), abductor pollicis brevis (APB), flexor pollicis brevis (FPB), extensor digitorum communis (EDC), flexor digitorum superficialis (FDS), extensor carpi radialis (ECR) and flexor carpi radialis (FCR) muscles.

### Experimental procedures

Participants were seated comfortably in an armchair, relaxing muscles of hands and arms. The experiment consisted of two identical sessions, *Session 1* and *Session 2*. These sessions were separated by one hour and served to test for test-retest reliability of our outcomes. EMG electrodes were kept fixed to minimise placement errors. The interval of one hour was set to prevent drying of the conductive electrolyte gel.

In each session, we searched for the hotspot positions for FDI, EDC and FCR before testing the RMTs. First, the stimulation intensity was identified that yielded MEPs for all three muscles when stimulating in the omega-shaped area (‘hand knob’) of the precentral gyrus. We started at 45% of the maximum stimulator output and increased or decreased the intensity until a consistent MEP was present. Then, we performed thirty stimulations around the hand knob region along the precentral gyrus. From these stimulations, we determined the position with the largest peak-to-peak amplitude for every muscle and labelled that position as the hotspot. Next, the RMT for every muscle was determined at the muscle-specific hotspot as the minimum stimulator output at which peak-to-peak amplitudes exceeded 50 μV in five out of ten stimuli. This was followed by the actual mapping procedure.

The TMS coil was pseudo-randomly positioned such that stimulations covered the entire left precentral gyrus. We applied 120 stimulations (Cavaleri et al., 2017) and repeated this at three intensities: 105% RMT of FDI, EDC and FCR, respectively. In total, we performed 360 stimulations in every session. We chose 105% RMT because previous studies suggested it to be the lowest possible intensity for upper limb muscles mapping (Krieg et al., 2017), thus leading to the least stimulation cross-talk. Finally, we estimated the hotspots of the other five muscles (ADM, APB, FPB, FDS and ECR) and determined the respective RMTs.

### MEP definition

We discriminated between TMS with and without eliciting MEPs, i.e., MEP and non-MEP points. MEPs were considered proper if their amplitude exceeded 20 times the EMG-baseline’s standard deviation (defined over 100 ms prior to each stimulation) but stayed below 10 mV (to exclude movement and cable artifacts), and if the peak’s latency fell within the range of 5 to 50 ms after stimulation. All other stimulations were marked as non-MEP points; see below under *Outcome measures* for further details.

### Area estimate

The area estimates were based on triangulated cortical surface meshes that we extracted using Freesurfer (https://surfer.nmr.mgh.harvard.edu/fswiki, version 7). We imported the meshes into Brainstorm (https://neuroimage.usc.edu/brainstorm/, version 3) to ease converting between world and subject-specific MRI coordinates. Next to the original meshes with about 230,000 to 340,000 vertices dependent on the participant, we also generated low-resolution version by downsampling the mesh to either 15,000 or 100,000 vertices. This enabled us to test for effects of surface resolution. In all cases, we assigned the Mindboggle anatomical atlas (https://mindboggle.info, version 6; see also Klein et al., 2017) to select left primary motor cortex. The subsequent area construction consisted of the following steps:

i. Stimulation points were projected to the triangulated cortical surface mesh yielding a set of vertices as illustrated in Figure 2 and further detailed under *Outcome measures*.

**Figure 2.**
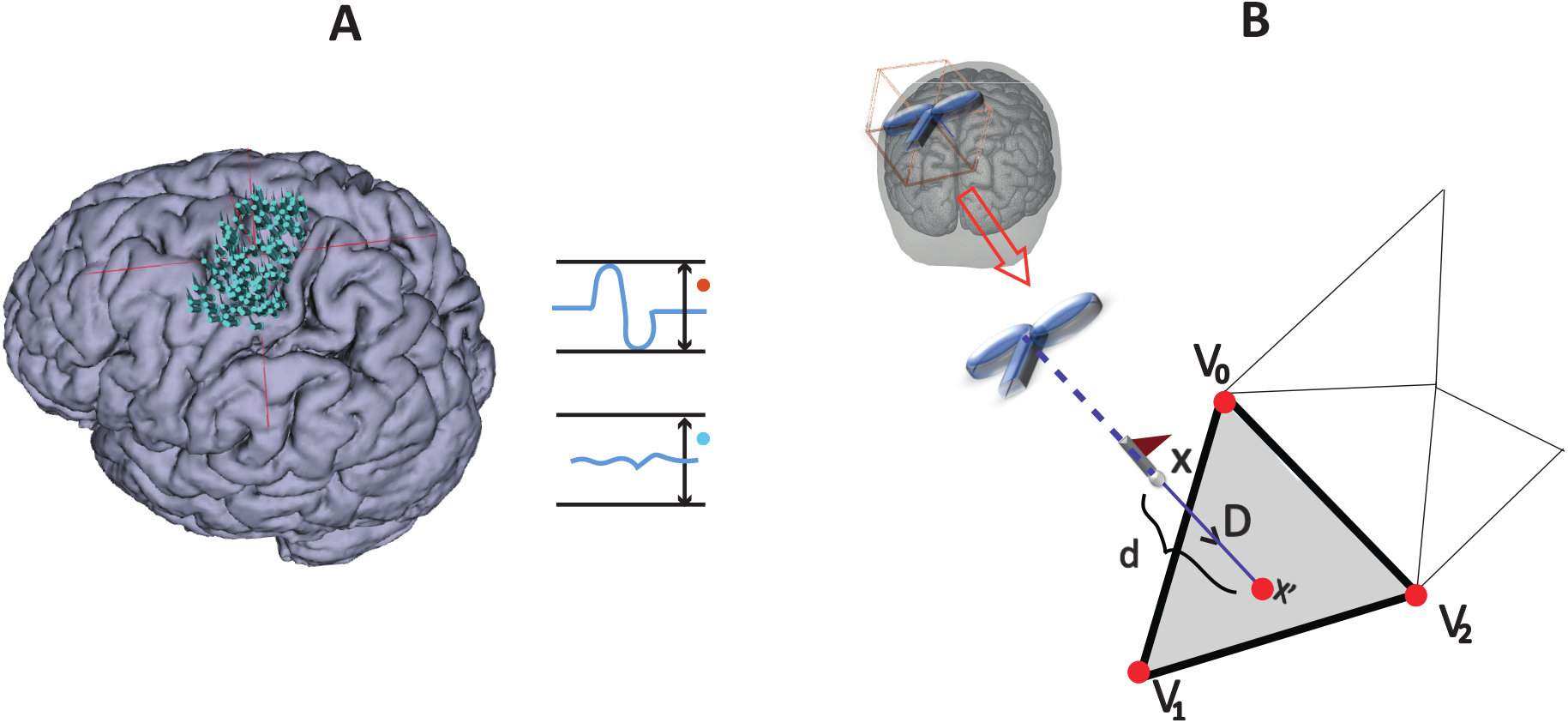
MEP definition and point projection along the direction of coil orientation. Panel A: Stimulation points over the cortex that either elicited an MEP or not: MEP and non-MEP points, respectively. Panel B: the cortical surface nearest to a given stimulation location along the TMS coil orientation is represented by the grey triangle (step ii). The three vertices in the triangle are shown by the red points, while the flagged spots indicate the stimulation points.
ii. Vertices that did not fall in left primary motor cortex were excluded (label ‘precentral L’ in the Mindboggle6 atlas).
iii. We connected the vertices along their shortest connecting paths. In brief, we converted the mesh into a sparse, weighted graph. The edges of triangularization served as adjacencies that we weighted by the Euclidean distance between the corresponding vertices. Then, we searched for the shortest paths between all MEP points (Dijkstra, 1959). We repeated this iteratively for all points of the connecting paths until no points were added. Figure 3 briefly summarises this iteration. Further details can be found at https://github.com/marlow17/surfaceanalysis.

**Figure 3.**
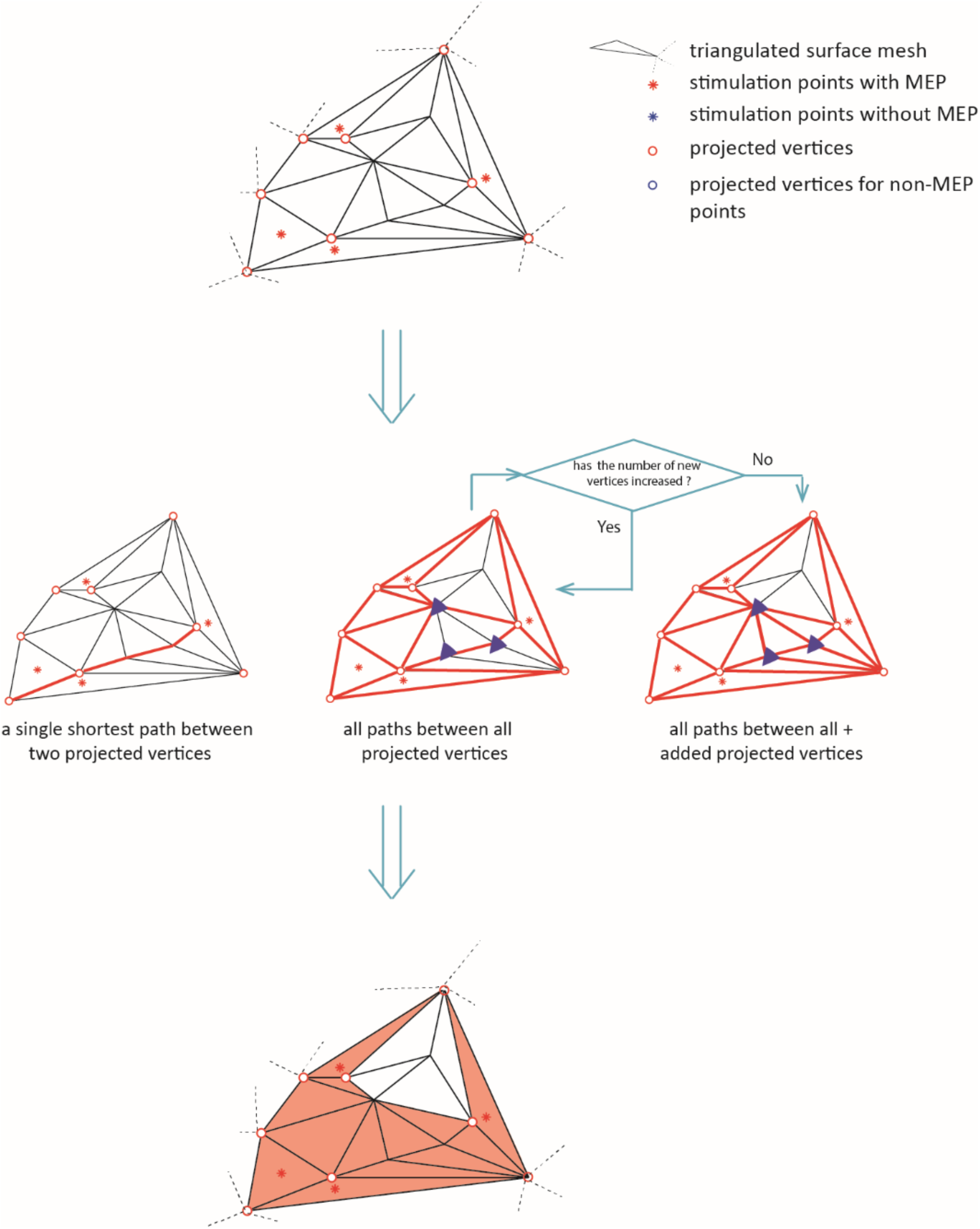
The diagram illustrates the iteration for defining the area of excitability (step iii). The red dots represent the active points and the blue dots the non-MEP one. Bold red lines indicate the shortest paths between the active points resulting in the orange shaded active area.
iv. Finally, we excluded the vertices (and triangles) corresponding to non-MEP points from the resulting area as sketched in Figure 4.

**Figure 4.**
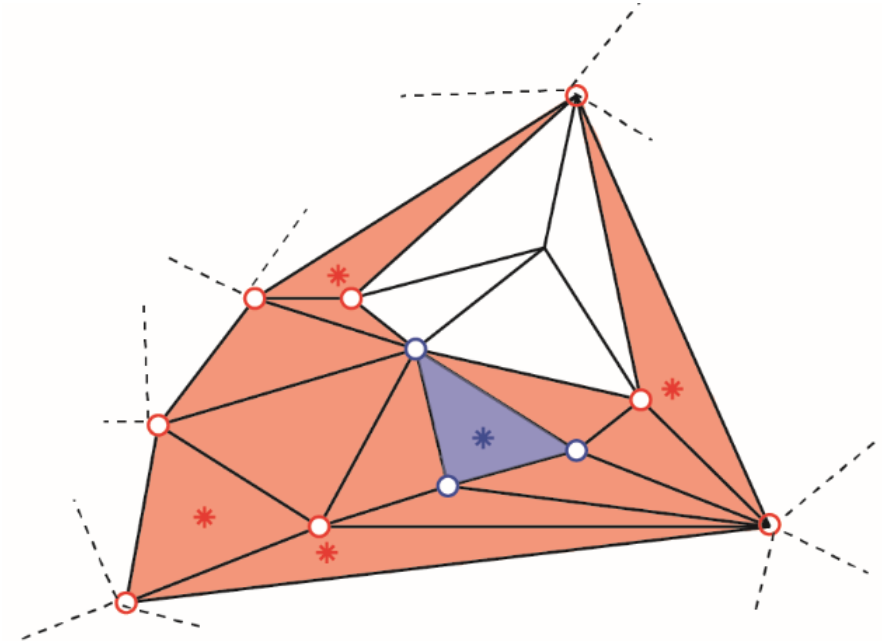
Removal of non-MEP points (step iv). The red dots represent the active points and the blue dots the non-MEP one. Bold lines indicate the shortest paths between the active points resulting in the orange shaded active area, and the blue background shows the (to-be-removed) non-active area.

Here we like to note that by construction, removing non-MEP points that are scattered across the area spanned by MEP points will reduce the size of the active areas. We briefly illustrate this in Figure 5.

**Figure 5.**
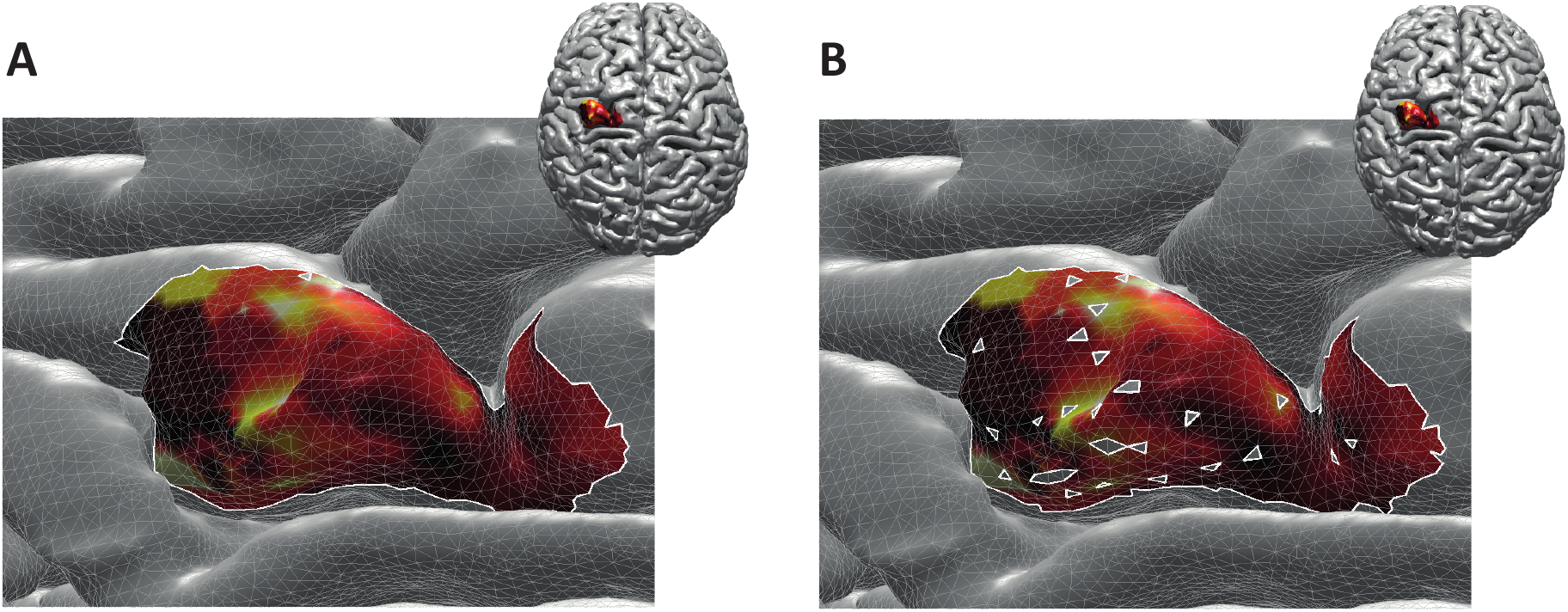
Reconstructed active area in primary motor cortex given a set of stimulation points (orange dots). The colour coding indicates the size of the MEP amplitude, with yellow being high and dark red being low. Panel A: Area without removing the non-MEP points (these points are marked in cyan); panel B: Area after removing the non-MEP points. By construction, the area shown in panel B is smaller than that in panel A. In both cases, the white lines represent an area’s boundary; note that this boundary does not necessarily equal the area’s convex hull, even when ignoring the non-MEP points (panel A).

From here on, we refer to the reconstruction without accounting for non-MEP points as method M1, whilst the removal of non-MEP points will be method M2 (with examples in Figure 5’s panels A and B, respectively).

### Outcome measures

Most of the outcome measures were based on the MEP amplitudes and latencies. We quantified them for every TMS pulse and for every muscle using the original EMG signals, from which we removed the stimulation artefact via linear interpolation (−1 to +2 ms around stimulation) followed by high pass filtering at 10 Hz (2^nd^ order, bi-directional Butterworth design). We defined the epoch −100 to −1 ms before the stimulus as a baseline and determined its mean value *μ* and the standard deviation *σ*. A peak in the interval 5 to 100 ms after the stimulus was considered an MEP if its value exceeded *μ* ± 20*σ*. Its latency was set as the first sample after the stimulus at which the signal exceeded *μ* + 2*σ* (*μ* − 2*σ* if the first peak is a maximum (minimum). Finally, we removed MEPs with peak-to-peak amplitude larger than 10 mV as we considered them artefacts.

The stimulation points were mapped onto cortical surfaces given as triangulated meshes. The triangle of the surface mesh closest to a stimulation point along the direction of coil orientation was determined following the approach by Möller and Trumbore (1997); see Figure 3. We assigned the vertices 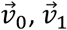, and 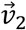 of the closest triangle the corresponding amplitude value, *a*. If two or more stimulations with an MEP shared a vertex, we averaged their amplitudes at the shared point. Since the total area of excitability possibly covered points that were not projected directly, we set all amplitude values via natural interpolation with C^1^ continuity to 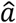; note that if interpolation was not needed, i.e., at the original vertices 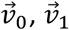, and 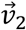, then 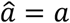. For every triangle we defined the length between their vertices as 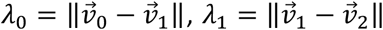 and 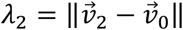, with ∥···∥ denoting the Euclidean distance between vertices. That is, given Cartesian coordinates 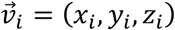, we used, e.g., 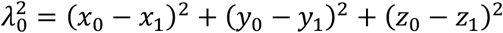.

The total area *A* of *k* = 1, …, *M* triangles weighted by the MEP amplitudes was computed via a slight modification of Heron’s formula, namely the triangular prism, that reads

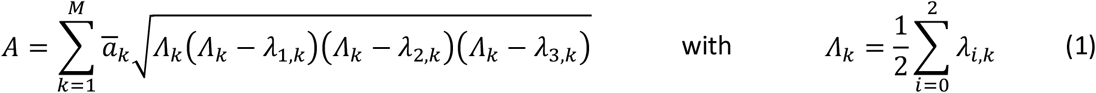

and 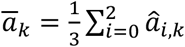 being the mean value of (interpolated) MEP amplitudes at the three vertices of triangle *k*.

Given all *i* = 1,…, *N* area vertices, we further defined the centroid of the total area in line with the conventional form of the centre of gravity (Opitz et al., 2014) as

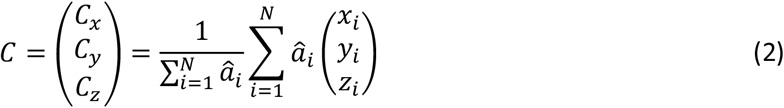

### Statistics

We first estimated the reliability between Session 1 and Session 2 via the intraclass correlation coefficients (ICC) of the centroid (*C*), the weighted area size (*A*) for every muscle and intensity level. We ran this analysis separately for the three representations of the cortex (i.e., three mesh resolutions) and for the three intensities. While the highest resolution for the intensity of 105% RMT of FCR will be reported below, all the other results can be found as *Supplementary Material*. There we also report the ICCs for centres of gravity (*CoG*) and for both the MEP amplitudes and the latencies.

To further confirm the absence of significant differences between Sessions 1 and 2, we performed a two-way ANOVA with repeated measures including factors of *intensity* and *session*. This also allowed for assessing effects of stimulation intensity. We applied a Bonferroni correction for multiple comparisons. Again, we restrict ourselves to reporting ‘only’ the findings of the estimates at maximum resolution in the body text and refer to the *Supplementary Material* for all other cases.

Finally, to assess effects of cortex mesh resolution and of ignoring/removing non-MEP stimulation points, we used a two-way repeated ANOVA with factors *method* and *resolution* (again with Bonferroni correction for multiple comparisons).

Prior to conducting the ANOVAs, sphericity was verified via Mauchly’s test. A Greenhouse-Geisser correction was performed if necessary. Throughout hypothesis testing, we used a significance threshold of α = .05. All statistical analyses were conducted using Matlab (The Mathworks Inc. Natwick, MA, version 2020b).

## Results

All *N* = 20 participants completed the experimental procedure without adverse reactions. Of all mappings (subjects × muscle × intensity × session = 960, each containing 120 stimulations) 2% did not contain any valid MEP, and thus did not enter further analyses. In 11/320 (subjects x muscles x session) cases this was for 105% RMT of FDI, 5/320 for 105% RMT of EDC, and 1/320 for 105% RMT of FCR; see *Supplementary* Table S1). For five subjects, we could not detect any MEPs for ADM when using the second intensity in both sessions.

When averaged over participants and sessions, the RMTs were FDI: 44.90±1.46%, ADM: 47.90±1.64%, APB: 46.15±1.45%, FPB: 46.78±1.73%, EDC: 45.28±1.50%, FDS: 47.75±1.51%, ECR: 46.55±1.50% and FCR: 48.00±1.52%, when expressed in stimulator intensity.

Table 1 provides an overview of ICC values obtained for maximum cortical resolution (results of the other resolutions and intensities can be found as *Supplementary* Tables S2a-c). For the sake of legibility, we defined distinct classes as follows: excellent: 0.8 ≤ ICC, good: 0.65 ≤ ICC < 0.8, moderate: 0.5 ≤ ICC < 0.65 and poor: ICC<0.5 (Cavaleri et al., 2018), and colour-coded the table entries accordingly.

**Table 1.** ICC values of area sizes *A* and centroids *C* = (*C_x_, C_y_, C_z_*)^*T*^ estimated for intensity of 105% RMT of FCR using the cortical meshes with maximum resolution when ignoring non-MEP points (M1) or removing them (M2).*

|  | FDI |  | ADM |  | APB |  | FPB |  | EDC |  | FDS |  | ECR |  | FCR |  |
| --- | --- | --- | --- | --- | --- | --- | --- | --- | --- | --- | --- | --- | --- | --- | --- | --- |
|  | M1 | M2 | M1 | M2 | M1 | M2 | M1 | M2 | M1 | M2 | M1 | M2 | M1 | M2 | M1 | M2 |
| $A$ | .59 | .59 | **.80** | **.81** | .71 | .70 | **.81** | **.81** | **.84** | **.84** | .49 | .47 | .74 | .73 | .25 | .24 |
| $C_x$ | **.95** | **.95** | **.94** | **.94** | **.94** | **.94** | **.94** | **.94** | **.96** | **.96** | **.94** | **.94** | **.96** | **.96** | **.95** | **.95** |
| $C_y$ | .72 | .73 | .70 | .70 | **.82** | **.82** | .75 | .76 | .71 | .71 | .65 | .65 | .66 | .65 | .69 | .69 |
| $C_z$ | **.89** | **.89** | **.81** | **.81** | **.86** | **.86** | **.90** | **.90** | **.86** | **.86** | .79 | .79 | **.84** | **.84** | **.91** | **.91** |

\* Excellent: $.8 \leq \text{ICC}$ (dark green, bold); good: $.65 \leq \text{ICC} < .8$ (light green); moderate: $.5 \leq \text{ICC} < .65$ (yellow); poor: $\text{ICC} < .5$ (light red).

The ICCs appeared consistent between methods M1 (ignoring non-MEP points) and M2 (removing non-MEP points). Most of them were good to excellent. This applies to the estimated centroids in the anterior/posterior and superior/inferior directions (*x* and *z* coordinates, respectively). The area sizes’ ICCs of FDI were moderate, and that of FDS and FCR were even poor. Yet, the ANOVA did not reveal any significant differences between the area estimates between sessions. We illustrate this in Table 2 for the highest cortex resolution and refer to the *supplementary* Tables S3a-b for the ANOVA results for the other cortex resolutions. In the *Supplementary* Tables S4/5/6a-c, we also provide the results for the corresponding centroid positions. In a nutshell there were hardly any significant effects of session or intensity (let alone their interaction) on the centroids; when correcting for multiple comparisons all effects will turn out not significant.

**Table 2.** Outcomes of the two-way ANOVA for the area sizes *A* (in mm^2^·μV·10^5^) with factors of *intensity* and *session* when considering the highest cortex mesh resolution and when removing the non-MEP points (M2).*

| | $\langle A \rangle$ at 105% RMT | | | *intensity* | | *session* | | *intensity* × *session* | | *p*-value pairwise comparison | | |
| --- | --- | --- | --- | --- | --- | --- | --- | --- | --- | --- | --- | --- |
|  | FDI | EDC | FCR | *F* | *p* | *F* | *p* | *F* | *p* | FDI/EDC | FDI/FCR | EDC/FCR |
| FDI | 1.61±0.24 | 2.48±0.60 | 4.86±1.41 | **F(2,36)=4.855** | **.032** | F(1,18)=1.390 | .254 | F(2,36)=0.634 | .454 | .307 | .087 | .195 |
| ADM | 0.92±0.21 | 0.99±0.17 | 2.16±0.60 | **F(2,24)=5.385** | **.032** | F(1,12)=1.216 | .292 | F(2,24)=0.238 | .790 | 1.00 | .076 | .137 |
| APB | 1.63±0.41 | 1.89±0.45 | 3.74±1.28 | F(2,34)=3.050 | .093 | F(1,17)=0.939 | .346 | F(2,34)=1.337 | .267 | 1.00 | .213 | .364 |
| FPB | 1.18±0.28 | 1.53±0.37 | 2.41±0.69 | F(2,34)=3.439 | .073 | F(1,17)=4.425 | .051 | F(2,34)=1.398 | .260 | .273 | .158 | .406 |
| EDC | 1.01±0.21 | 0.94±0.12 | 1.75±0.36 | **F(2,34)=5.960** | **.016** | F(1,17)=0.082 | .778 | F(2,34)=0.146 | .865 | 1.00 | **.031** | .078 |
| FDS | 0.83±0.12 | 0.97±0.18 | 1.60±0.25 | **F(2,30)=6.345** | **.005** | F(1,15)=1.090 | .313 | F(2,30)=1.174 | .323 | 1.00 | **.020** | .077 |
| ECR | 1.24±0.20 | 1.62±0.43 | 2.45±0.53 | **F(2,30)=4.730** | **.016** | F(1,15)=0.319 | .580 | F(2,30)=0.867 | .388 | .947 | **.030** | .217 |
| FCR | 0.69±0.11 | 0.99±0.20 | 1.39±0.20 | **F(2,30)=5.173** | **.012** | F(1,15)=2.079 | .170 | F(2,30)=1.388 | .265 | .473 | **.011** | .376 |

\* Bold face implies $p < .05$ .

Here we would like to note that this dependency on stimulation intensity can be understood when looking at the effects of intensity on the mere MEP amplitudes, i.e., without projecting them onto the cortex. The corresponding results can be found as *Supplementary* Table S7. In a nutshell, the amplitudes of FDI, EDC, FCR, FDS and FCR significantly increased with increasing stimulation intensity.

As expected, ignoring non-MEP points (M1) consistently resulted in larger area sizes when compared to the case when non-MEP points were removed (M2). Our second ANOVA confirmed this. We summarized this in Table 3 where we highlighted the main effects of *method*. Yet, we also would like to note the interaction effect with *resolution*, suggesting that the correction for non-MEP points is especially relevant when incorporating low-resolution cortical meshes (see also Figure 7, upper row).

**Table 3.**
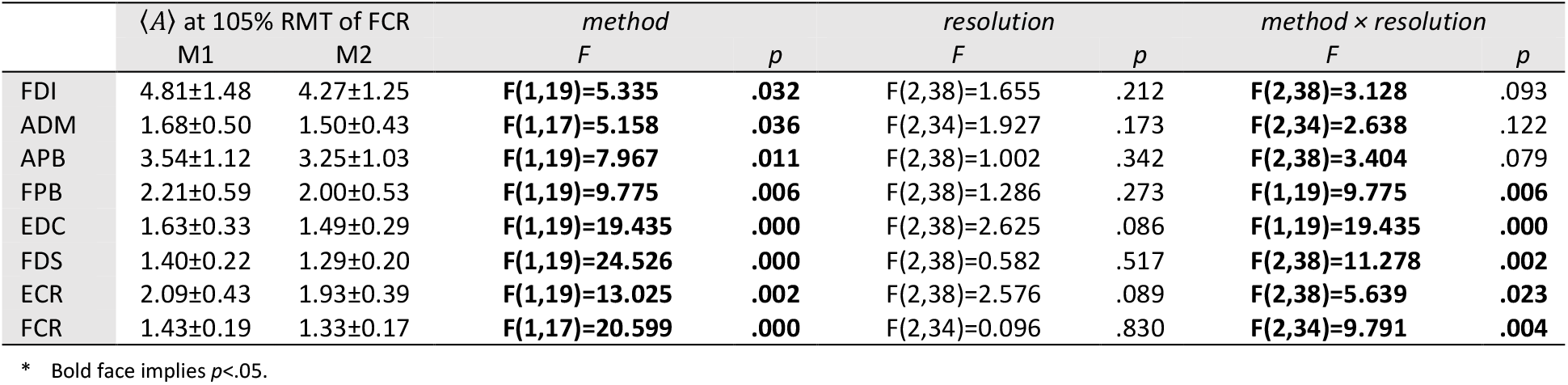
The outcome of the two-way ANOVA for the area sizes *A* (in mm^2^·μV·10^5^) with factors of *method* and *resolution* for the intensity of 105% RMT of FCR; M1 = ignoring non-MEP points, M2 = removing them.*

| | $\langle A \rangle$ at 105% RMT of FCR | | *method* | | *resolution* | | *method × resolution* | |
| --- | --- | --- | --- | --- | --- | --- | --- | --- |
|  | M1 | M2 | *F* | *p* | *F* | *p* | *F* | *p* |
| FDI | 4.81±1.48 | 4.27±1.25 | **F(1,19)=5.335** | **.032** | F(2,38)=1.655 | .212 | **F(2,38)=3.128** | .093 |
| ADM | 1.68±0.50 | 1.50±0.43 | **F(1,17)=5.158** | **.036** | F(2,34)=1.927 | .173 | **F(2,34)=2.638** | .122 |
| APB | 3.54±1.12 | 3.25±1.03 | **F(1,19)=7.967** | **.011** | F(2,38)=1.002 | .342 | **F(2,38)=3.404** | .079 |
| FPB | 2.21±0.59 | 2.00±0.53 | **F(1,19)=9.775** | **.006** | F(2,38)=1.286 | .273 | **F(1,19)=9.775** | **.006** |
| EDC | 1.63±0.33 | 1.49±0.29 | **F(1,19)=19.435** | **.000** | F(2,38)=2.625 | .086 | **F(1,19)=19.435** | **.000** |
| FDS | 1.40±0.22 | 1.29±0.20 | **F(1,19)=24.526** | **.000** | F(2,38)=0.582 | .517 | **F(2,38)=11.278** | **.002** |
| ECR | 2.09±0.43 | 1.93±0.39 | **F(1,19)=13.025** | **.002** | F(2,38)=2.576 | .089 | **F(2,38)=5.639** | **.023** |
| FCR | 1.43±0.19 | 1.33±0.17 | **F(1,17)=20.599** | **.000** | F(2,34)=0.096 | .830 | **F(2,34)=9.791** | **.004** |

\* Bold face implies $p < .05$ .

The main effect of *method* (ignoring non-MEPs versus removing them) is also illustrated in Figure 6 where we show the relative change in the estimated area sizes. Irrespective of resolution, not removing the non-MEP points yields an overestimation of the active areas, though this effect appears particularly pronounced at low resolution (top panel in Figure 6).

**Figure 6.**
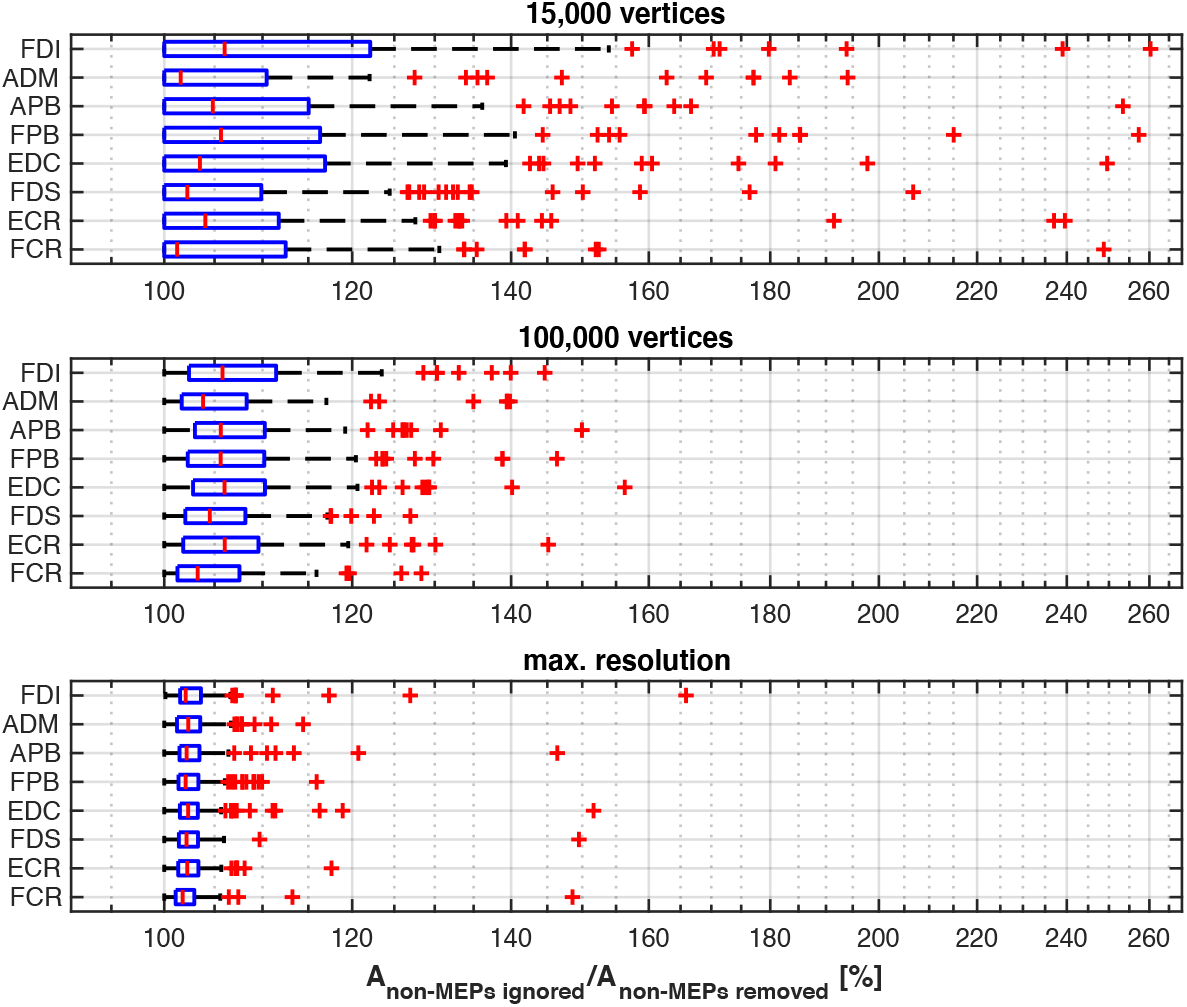
The relative change of the size of the estimated active area per muscle using different resolutions of the cortex mesh. The maximum number of vertices was subject specific and ranged from about 230,000 to 340,000 vertices. The figure shows boxplots with red crosses marking outliers (all subjects/sessions entered the median and quantile estimates). Throughout mesh resolution the removal of non-MEP points yielded larger area sizes suggesting an overestimation of the active area.

We finally illustrate the effect of removing non-MEP points in Figure 7, where it can be clearly seen that higher resolutions lead to less area being removed.

**Figure 7.**
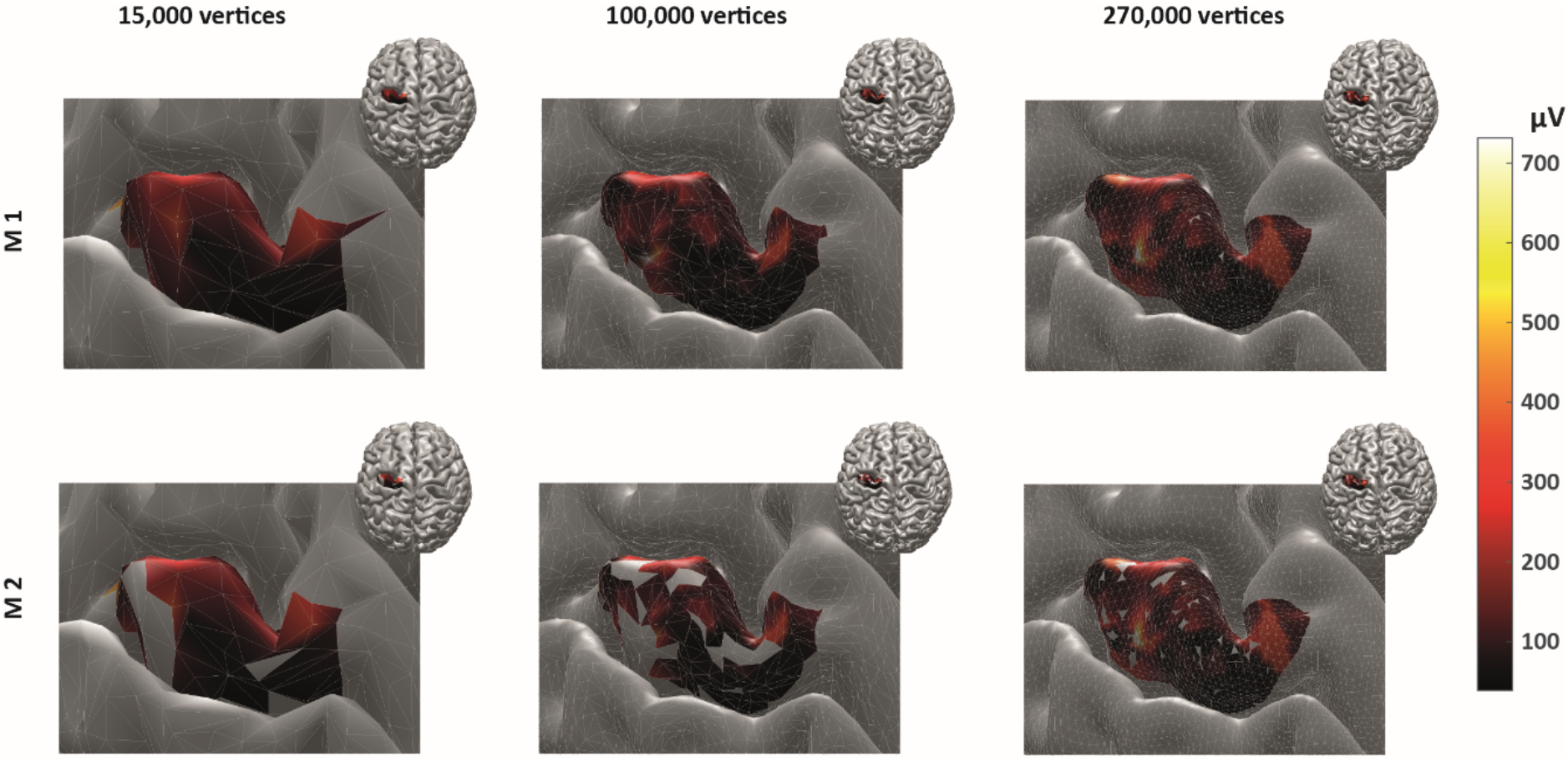
The cortical representation for Session 1 intensity of 105% RMT of FCR in a representative participant. In M1 (upper row), we ignored non-MEP points, and in M2 (lower row), we removed them. Left: surface mesh resolution of 15,000 vertices; centre: 100,000 vertices; right: maximum number of vertex number (in this participant, about 270,000 vertices). Colour coding represents the value of the (interpolated) MEP-amplitude (in μV). The effects of non-MEP point removal are especially visible at a lower resolution, but in all cases, removing these stimulation points cause the estimated area to shrink.

## Discussion

We assessed the reliability of the cortical representation of eight muscles mapped simultaneously using navigated TMS. We distinguished two methods to estimate the active area of a muscle. In the first, more conventional one (M1), we included all stimulation points that elicited an MEP. In the second method (M2), we included the same points but also excluded all stimulation points that did not elicit an MEP. We tested for the effects of the type of measure with the obvious expectation that the latter will yield smaller active areas. We also tested for effects of stimulation intensity and cortical mesh resolution in two consecutive sessions. We found that the reliabilities of the size and the centroids of the active areas for all the muscles were mostly good or excellent. Exceptions were the area size estimates in three muscles (Table 1) that came with small areas sizes but strong outliers when looking at their representation at high-resolution cortical surface meshes (Figure 6, lower panel). Note that the reliability of amplitude and latency were excellent or good for all the muscles, and all the outcomes analysed in ECR achieved excellent ICCs (see *Supplementary Material* S8), indicating the reliability of our experimental approach.

One must realise that designing multiple muscle mapping experiments can – in general – be problematic as the RMT of a single muscle must be considered a reference when setting the stimulation intensity. In our findings, the values of average RMTs indicated that the difference between RMTs is small (not more than 3.1% of stimulator output). Intensities of 105%, 110% to 120% (Akiyama et al., 2006) RMT have been widely used in motor mapping (Akiyama et al., 2006; Bohning et al., 2001; Tarapore et al., 2012), suggesting that the here-observed difference is acceptable if not negligible. Hence, forearm and hand muscles might be pooled in a group of muscles with “similar RMTs” and may indeed be evaluated at the same intensity.

The ICCs of centroids for all the muscles were excellent or good (see *Supplementary Material* S2a-c for the likewise good results for the more conventional CoGs). Put differently, the estimated centres-of-gravity appeared very consistent and should be considered a reliable outcome in motor mapping, in line with previous studies (Cavaleri et al., 2018; Weiss et al., 2013). The CoG is commonly employed to quantify the cortical representation of muscles (Massé-Alarie et al., 2017; Nazarova et al., 2021). However, there are several issues with the notion of ‘CoG’ itself. For instance, for many shapes (of cortical representations), the CoG will lie outside the actual stimulation area itself (consider a banana, whose CoG will not be inside the banana itself). The CoG may hence be a tricky measure to give an estimate of the cortical representation of a muscle, especially when the true cortical representation is non-trivially shaped and on a curved surface. Supplementing the CoG, or in our case the centroid, by the area of excitability is clearly needed, especially when the area estimate is weighted by the MEP amplitude. Again, we advocate incorporating the non-MEP stimulation points in these estimates.

When removing non-MEP points, the areas of excitability became significantly smaller than when non-MEP points were simply ignored. We argue that by ignoring the stimulation points that do not elicit MEPs one runs the risk of overestimating the area of excitability and thus to mis-represent muscles in the cortex. One may counter this idea by subsuming that the neuronal population that ought to be covered by our cortical map are likely to be homogeneously distributed. However, several invasive studies already speak against this (e.g., Schieber, 2001). By using intracortical micro-stimulation, Nudo et al. (1996) revealed that the cortical representation of distal forelimb muscles is quite complicated and clearly not uniform.

## Conclusion

Estimating the active area can be improved when incorporation points at which TMS does not elicit an MEP. Navigated TMS and a pseudo-random coil placement allow for correcting area estimates post-hoc and hence reduce the risk of overestimating the cortical representation of active areas. As such, the very fact that at certain points, a stimulation does not yield a measurable response appears informative. And, even when assessing multiple muscles in unison, this approach comes with high reliability, albeit under the proviso that stimulation intensity has been chosen properly.

