## Supplemental Table s1-s8 for "Accounting for stimulations that do not elicit motor-evoked potentials when mapping cortical representations of multiple muscles"

### *Stimulations that elicited MEPs*

**Table S1**. Overview of elicited MEPs in all the 20 subjects × 8 muscles × 3 intensities × 2 sessions=960 mappings; 'A' stands for active, and' N' stands for non-active.

| | Intensity of 105% RMT of FDI | | | | | | | | | | | | | | | |
|---|---|---|---|---|---|---|---|---|---|---|---|---|---|---|---|---|
| | Session1 | | | | | | | | Session2 | | | | | | | |
| | FDI | ADM | APB | FPB | EDC | FDS | ECR | FCR | FDI | ADM | APB | FPB | EDC | FDS | ECR | FCR |
| Subject01 | 'A' | 'A' | 'A' | 'A' | 'A' | 'A' | 'A' | 'A' | 'A' | 'A' | 'A' | 'A' | 'A' | 'A' | 'A' | 'A' |
| Subject02 | 'A' | 'A' | 'A' | 'A' | 'A' | 'A' | 'A' | 'A' | 'A' | 'A' | 'A' | 'A' | 'A' | 'A' | 'A' | 'A' |
| Subject03 | 'A' | 'A' | 'A' | 'A' | 'A' | 'A' | 'A' | 'A' | 'A' | 'A' | 'A' | 'A' | 'A' | 'A' | 'A' | 'A' |
| Subject04 | 'A' | 'A' | 'A' | 'A' | 'A' | 'A' | 'A' | 'A' | 'A' | 'A' | 'A' | 'A' | 'A' | 'A' | 'A' | 'A' |
| Subject05 | 'A' | 'A' | 'A' | 'A' | 'A' | 'A' | 'A' | 'A' | 'A' | 'A' | 'A' | 'A' | 'A' | 'A' | 'A' | 'A' |
| Subject06 | 'A' | 'A' | 'A' | 'A' | 'A' | 'A' | 'A' | 'A' | 'A' | 'A' | 'A' | 'A' | 'A' | 'A' | 'A' | 'A' |
| Subject07 | 'A' | 'A' | 'A' | 'A' | 'A' | 'A' | 'A' | 'A' | 'A' | 'A' | 'A' | 'A' | 'A' | 'A' | 'A' | 'A' |
| Subject08 | 'A' | 'A' | 'A' | 'A' | 'A' | 'A' | 'A' | 'A' | 'A' | 'A' | 'A' | 'A' | 'A' | 'A' | 'A' | 'A' |
| Subject09 | 'A' | 'A' | 'A' | 'A' | 'A' | 'A' | 'A' | 'A' | 'A' | 'A' | 'A' | 'A' | 'A' | 'A' | 'A' | 'A' |
| Subject10 | 'A' | 'A' | 'A' | 'A' | 'A' | 'A' | 'A' | 'A' | 'A' | 'A' | 'A' | 'A' | 'A' | 'A' | 'A' | 'A' |
| Subject11 | 'A' | 'A' | 'A' | 'A' | 'A' | 'A' | 'A' | 'A' | 'A' | 'A' | 'A' | 'A' | 'A' | 'A' | 'A' | 'A' |
| Subject12 | 'A' | 'A' | 'A' | 'A' | 'A' | 'A' | 'A' | 'A' | 'A' | 'A' | 'A' | 'A' | 'A' | 'A' | 'A' | 'A' |
| Subject13 | 'A' | 'A' | 'A' | 'A' | 'A' | 'A' | 'A' | 'A' | 'A' | 'A' | 'A' | 'A' | 'A' | 'A' | 'A' | 'A' |
| Subject14 | 'A' | 'A' | 'A' | 'A' | 'A' | 'A' | 'A' | 'A' | 'A' | 'N' | 'A' | 'A' | 'A' | 'A' | 'A' | 'A' |
| Subject15 | 'A' | 'A' | 'A' | 'A' | 'A' | 'A' | 'A' | 'A' | 'A' | 'A' | 'A' | 'A' | 'A' | 'A' | 'A' | 'A' |
| Subject16 | 'A' | 'A' | 'A' | 'A' | 'A' | 'A' | 'A' | 'A' | 'A' | 'A' | 'A' | 'A' | 'A' | 'A' | 'A' | 'A' |
| Subject17 | 'A' | 'A' | 'A' | 'A' | 'A' | 'A' | 'A' | 'A' | 'A' | 'A' | 'A' | 'A' | 'A' | 'A' | 'A' | 'A' |
| Subject18 | 'A' | 'N' | 'A' | 'A' | 'A' | 'N' | 'A' | 'A' | 'A' | 'A' | 'A' | 'A' | 'A' | 'A' | 'A' | 'A' |
| Subject19 | 'A' | 'A' | 'A' | 'A' | 'A' | 'A' | 'A' | 'A' | 'A' | 'A' | 'A' | 'A' | 'A' | 'A' | 'A' | 'A' |
| Subject20 | 'N' | 'N' | 'N' | 'N' | 'N' | 'N' | 'N' | 'N' | 'A' | 'A' | 'A' | 'A' | 'A' | 'A' | 'A' | 'A' |

| | Intensity of 105% RMT of EDC | | | | | | | | | | | | | | | |
|---|---|---|---|---|---|---|---|---|---|---|---|---|---|---|---|---|
| | Session1 | | | | | | | | Session2 | | | | | | | |
| | FDI | ADM | APB | FPB | EDC | FDS | ECR | FCR | FDI | ADM | APB | FPB | EDC | FDS | ECR | FCR |
| Subject01 | 'A' | 'A' | 'A' | 'A' | 'A' | 'A' | 'A' | 'A' | 'A' | 'A' | 'A' | 'A' | 'A' | 'A' | 'A' | 'A' |
| Subject02 | 'A' | 'A' | 'A' | 'A' | 'A' | 'A' | 'A' | 'A' | 'A' | 'A' | 'A' | 'A' | 'A' | 'A' | 'A' | 'A' |
| Subject03 | 'A' | 'A' | 'A' | 'A' | 'A' | 'A' | 'A' | 'A' | 'A' | 'A' | 'A' | 'A' | 'A' | 'A' | 'A' | 'A' |
| Subject04 | 'A' | 'A' | 'A' | 'A' | 'A' | 'A' | 'A' | 'A' | 'A' | 'A' | 'A' | 'A' | 'A' | 'A' | 'A' | 'A' |
| Subject05 | 'A' | 'A' | 'A' | 'A' | 'A' | 'A' | 'A' | 'A' | 'A' | 'A' | 'A' | 'A' | 'A' | 'A' | 'A' | 'A' |
| Subject06 | 'A' | 'A' | 'A' | 'A' | 'A' | 'A' | 'A' | 'A' | 'A' | 'A' | 'A' | 'A' | 'A' | 'A' | 'A' | 'A' |
| Subject07 | 'A' | 'A' | 'A' | 'A' | 'A' | 'A' | 'A' | 'A' | 'A' | 'A' | 'A' | 'A' | 'A' | 'A' | 'A' | 'A' |
| Subject08 | 'A' | 'A' | 'A' | 'A' | 'A' | 'A' | 'A' | 'A' | 'A' | 'A' | 'A' | 'A' | 'A' | 'A' | 'A' | 'A' |
| Subject09 | 'A' | 'A' | 'A' | 'A' | 'A' | 'A' | 'A' | 'A' | 'A' | 'A' | 'A' | 'A' | 'A' | 'A' | 'A' | 'A' |
| Subject10 | 'A' | 'A' | 'A' | 'A' | 'A' | 'A' | 'A' | 'A' | 'A' | 'A' | 'A' | 'A' | 'A' | 'A' | 'A' | 'A' |
| Subject11 | 'A' | 'A' | 'A' | 'A' | 'A' | 'A' | 'A' | 'N' | 'A' | 'A' | 'A' | 'A' | 'A' | 'A' | 'A' | 'A' |
| Subject12 | 'A' | 'N' | 'A' | 'A' | 'A' | 'A' | 'A' | 'A' | 'A' | 'N' | 'A' | 'A' | 'A' | 'A' | 'A' | 'A' |
| Subject13 | 'A' | 'A' | 'A' | 'A' | 'A' | 'A' | 'A' | 'A' | 'A' | 'A' | 'A' | 'A' | 'A' | 'A' | 'A' | 'A' |
| Subject14 | 'A' | 'N' | 'A' | 'A' | 'A' | 'A' | 'A' | 'A' | 'A' | 'N' | 'A' | 'A' | 'A' | 'A' | 'A' | 'A' |
| Subject15 | 'A' | 'A' | 'A' | 'A' | 'A' | 'A' | 'A' | 'A' | 'A' | 'A' | 'A' | 'A' | 'A' | 'A' | 'A' | 'A' |
| Subject16 | 'A' | 'A' | 'A' | 'A' | 'A' | 'A' | 'A' | 'A' | 'A' | 'A' | 'A' | 'A' | 'A' | 'A' | 'A' | 'A' |
| Subject17 | 'A' | 'A' | 'A' | 'A' | 'A' | 'A' | 'A' | 'A' | 'A' | 'A' | 'A' | 'A' | 'A' | 'A' | 'A' | 'A' |
| Subject18 | 'A' | 'A' | 'A' | 'A' | 'A' | 'A' | 'A' | 'A' | 'A' | 'A' | 'A' | 'A' | 'A' | 'A' | 'A' | 'A' |
| Subject19 | 'A' | 'A' | 'A' | 'A' | 'A' | 'A' | 'A' | 'A' | 'A' | 'A' | 'A' | 'A' | 'A' | 'A' | 'A' | 'A' |
| Subject20 | 'A' | 'A' | 'A' | 'A' | 'A' | 'A' | 'A' | 'A' | 'A' | 'A' | 'A' | 'A' | 'A' | 'A' | 'A' | 'A' |

| | Intensity of 105% RMT of FCR | | | | | | | | | | | | | | | |
|---|---|---|---|---|---|---|---|---|---|---|---|---|---|---|---|---|
| | Session1 | | | | | | | | Session2 | | | | | | | |
| | FDI | ADM | APB | FPB | EDC | FDS | ECR | FCR | FDI | ADM | APB | FPB | EDC | FDS | ECR | FCR |
| Subject01 | 'A' | 'A' | 'A' | 'A' | 'A' | 'A' | 'A' | 'A' | 'A' | 'A' | 'A' | 'A' | 'A' | 'A' | 'A' | 'A' |
| Subject02 | 'A' | 'A' | 'A' | 'A' | 'A' | 'A' | 'A' | 'A' | 'A' | 'A' | 'A' | 'A' | 'A' | 'A' | 'A' | 'A' |
| Subject03 | 'A' | 'A' | 'A' | 'A' | 'A' | 'A' | 'A' | 'A' | 'A' | 'A' | 'A' | 'A' | 'A' | 'A' | 'A' | 'A' |
| Subject04 | 'A' | 'A' | 'A' | 'A' | 'A' | 'A' | 'A' | 'A' | 'A' | 'A' | 'A' | 'A' | 'A' | 'A' | 'A' | 'A' |
| Subject05 | 'A' | 'A' | 'A' | 'A' | 'A' | 'A' | 'A' | 'A' | 'A' | 'A' | 'A' | 'A' | 'A' | 'A' | 'A' | 'A' |
| Subject06 | 'A' | 'A' | 'A' | 'A' | 'A' | 'A' | 'A' | 'A' | 'A' | 'A' | 'A' | 'A' | 'A' | 'A' | 'A' | 'A' |
| Subject07 | 'A' | 'A' | 'A' | 'A' | 'A' | 'A' | 'A' | 'A' | 'A' | 'A' | 'A' | 'A' | 'A' | 'A' | 'A' | 'A' |
| Subject08 | 'A' | 'A' | 'A' | 'A' | 'A' | 'A' | 'A' | 'A' | 'A' | 'A' | 'A' | 'A' | 'A' | 'A' | 'A' | 'A' |

| | | | | | | | | | | | | | | | | |
|---|---|---|---|---|---|---|---|---|---|---|---|---|---|---|---|---|
| **Subject09** | 'A' | 'A' | 'A' | 'A' | 'A' | 'A' | 'A' | 'A' | 'A' | 'A' | 'A' | 'A' | 'A' | 'A' | 'A' | 'A' |
| **Subject10** | 'A' | 'A' | 'A' | 'A' | 'A' | 'A' | 'A' | 'A' | 'A' | 'A' | 'A' | 'A' | 'A' | 'A' | 'A' | 'A' |
| **Subject11** | 'A' | 'A' | 'A' | 'A' | 'A' | 'A' | 'A' | 'A' | 'A' | 'A' | 'A' | 'A' | 'A' | 'A' | 'A' | 'A' |
| **Subject12** | 'A' | 'N' | 'A' | 'A' | 'A' | 'A' | 'A' | 'A' | 'A' | 'A' | 'A' | 'A' | 'A' | 'A' | 'A' | 'A' |
| **Subject13** | 'A' | 'A' | 'A' | 'A' | 'A' | 'A' | 'A' | 'A' | 'A' | 'A' | 'A' | 'A' | 'A' | 'A' | 'A' | 'A' |
| **Subject14** | 'A' | 'A' | 'A' | 'A' | 'A' | 'A' | 'A' | 'A' | 'A' | 'A' | 'A' | 'A' | 'A' | 'A' | 'A' | 'A' |
| **Subject15** | 'A' | 'A' | 'A' | 'A' | 'A' | 'A' | 'A' | 'A' | 'A' | 'A' | 'A' | 'A' | 'A' | 'A' | 'A' | 'A' |
| **Subject16** | 'A' | 'A' | 'A' | 'A' | 'A' | 'A' | 'A' | 'A' | 'A' | 'A' | 'A' | 'A' | 'A' | 'A' | 'A' | 'A' |
| **Subject17** | 'A' | 'A' | 'A' | 'A' | 'A' | 'A' | 'A' | 'A' | 'A' | 'A' | 'A' | 'A' | 'A' | 'A' | 'A' | 'A' |
| **Subject18** | 'A' | 'A' | 'A' | 'A' | 'A' | 'A' | 'A' | 'A' | 'A' | 'A' | 'A' | 'A' | 'A' | 'A' | 'A' | 'A' |
| **Subject19** | 'A' | 'A' | 'A' | 'A' | 'A' | 'A' | 'A' | 'A' | 'A' | 'A' | 'A' | 'A' | 'A' | 'A' | 'A' | 'A' |
| **Subject20** | 'A' | 'A' | 'A' | 'A' | 'A' | 'A' | 'A' | 'A' | 'A' | 'A' | 'A' | 'A' | 'A' | 'A' | 'A' | 'A' |

### *Number of excitable (active) points*

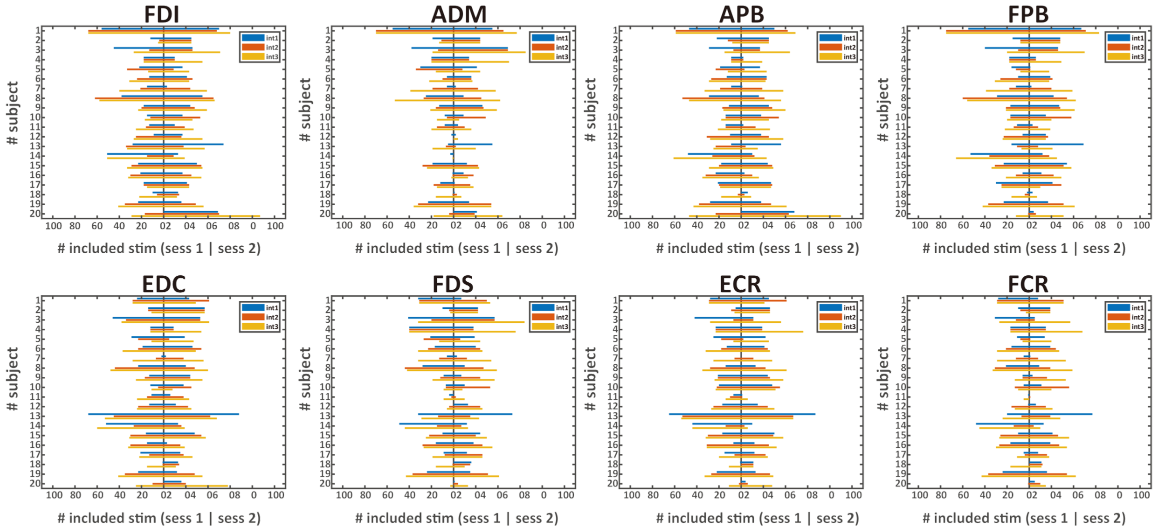


**Figure S1**. The figure depicts the number of active points for all the muscles in all the subjects. The x-axis values show the number of stimulation of session 1 (centre to the left) to session 2 (centre to the right). Y-axis values represent the number of subjects. The blue, red, and yellow colour legends show the intensity 1,2 and 3 values, respectively.

### *Supplementary statistics*

**Table S2a**. ICC values of area sizes $A$ and centroids $C = \left(C_x, C_y, C_z\right)^T$ estimated for intensity of 105% RMT of FCR using the cortical meshes with maximum resolution when ignoring non-MEP points (M1) or removing them (M2).*

| | FDI | | ADM | | APB | | FPB | | EDC | | FDS | | ECR | | FCR | |
|---|---|---|---|---|---|---|---|---|---|---|---|---|---|---|---|---|
| | M1 | M2 | M1 | M2 | M1 | M2 | M1 | M2 | M1 | M2 | M1 | M2 | M1 | M2 | M1 | M2 |
| Resolution: 15,000 vertices | | | | | | | | | | | | | | | | |
| $A$ | 0.44 | 0.60 | 0.79 | **0.86** | **0.80** | 0.61 | **0.92** | 0.74 | **0.87** | **0.84** | 0.49 | 0.46 | **0.84** | 0.76 | 0.40 | 0.38 |
| $C_x$ | **0.94** | **0.94** | **0.94** | **0.94** | **0.94** | **0.93** | **0.95** | **0.95** | **0.96** | **0.95** | **0.95** | **0.94** | **0.97** | **0.96** | **0.96** | **0.96** |
| $C_y$ | 0.76 | 0.73 | 0.68 | 0.68 | 0.75 | 0.73 | 0.73 | 0.70 | 0.75 | 0.73 | 0.70 | 0.68 | 0.75 | 0.71 | 0.75 | 0.67 |
| $C_z$ | **0.90** | **0.89** | **0.84** | **0.83** | **0.85** | **0.84** | **0.88** | **0.87** | **0.89** | **0.89** | **0.83** | **0.82** | **0.89** | **0.88** | **0.90** | **0.87** |
| Resolution: 100,000 vertices | | | | | | | | | | | | | | | | |
| $A$ | 0.54 | 0.57 | **0.84** | **0.85** | 0.72 | 0.71 | **0.87** | **0.85** | **0.91** | **0.91** | 0.66 | 0.64 | **0.87** | **0.87** | 0.46 | 0.43 |
| $C_x$ | **0.95** | **0.95** | **0.94** | **0.94** | **0.94** | **0.94** | **0.94** | **0.93** | **0.95** | **0.95** | **0.93** | **0.94** | **0.96** | **0.96** | **0.95** | **0.95** |
| $C_y$ | 0.71 | 0.71 | 0.69 | 0.69 | **0.80** | **0.80** | 0.76 | 0.76 | 0.75 | 0.74 | 0.66 | 0.64 | 0.67 | 0.66 | 0.71 | 0.70 |
| $C_z$ | **0.90** | **0.90** | **0.81** | **0.81** | **0.87** | **0.88** | **0.89** | **0.89** | **0.88** | **0.88** | **0.80** | **0.80** | **0.88** | **0.88** | **0.92** | **0.92** |

* Excellent: .8 ≤ ICC (dark green, bold); good: .65 ≤ ICC < .8 (light green); moderate: .5 ≤ ICC < .65 (yellow); poor: ICC<.5 (light red).

**Table S2b**. ICC values of area sizes $A$ and centroids $C = (C_x, C_y, C_z)^T$ estimated for intensity of 105% RMT of FDI using the cortical meshes with maximum resolution when ignoring non-MEP points (M1) or removing them (M2).*

| | FDI | | ADM | | APB | | FPB | | EDC | | FDS | | ECR | | FCR | |
|---|---|---|---|---|---|---|---|---|---|---|---|---|---|---|---|---|
| | M1 | M2 | M1 | M2 | M1 | M2 | M1 | M2 | M1 | M2 | M1 | M2 | M1 | M2 | M1 | M2 |
| Resolution: 15,000 vertices | | | | | | | | | | | | | | | | |
| $A$ | 0.78 | 0.78 | 0.44 | 0.50 | **0.83** | **0.85** | **0.55** | 0.54 | 0.65 | 0.31 | 0.68 | 0.70 | 0.75 | 0.63 | 0.63 | 0.61 |
| $C_x$ | **0.96** | **0.95** | **0.94** | **0.95** | **0.93** | **0.93** | **0.93** | **0.92** | **0.98** | **0.98** | **0.97** | **0.96** | **0.97** | **0.97** | **0.97** | **0.96** |
| $C_y$ | **0.80** | 0.79 | 0.68 | 0.66 | **0.81** | **0.82** | **0.82** | **0.82** | 0.78 | 0.78 | 0.78 | 0.79 | 0.78 | 0.77 | **0.81** | **0.81** |
| $C_z$ | **0.90** | **0.89** | **0.89** | **0.88** | **0.85** | **0.84** | **0.83** | **0.83** | **0.88** | **0.87** | **0.90** | **0.90** | **0.90** | **0.88** | **0.85** | **0.85** |
| Resolution: 100,000 vertices | | | | | | | | | | | | | | | | |
| $A$ | 0.73 | 0.72 | 0.19 | 0.16 | **0.83** | **0.83** | **0.58** | 0.59 | 0.58 | 0.50 | 0.65 | 0.68 | 0.78 | 0.71 | 0.56 | 0.55 |
| $C_x$ | **0.94** | **0.94** | **0.92** | **0.92** | **0.92** | **0.92** | **0.92** | **0.92** | **0.98** | **0.98** | **0.95** | **0.95** | **0.97** | **0.97** | **0.94** | **0.94** |
| $C_y$ | **0.85** | **0.86** | 0.76 | 0.72 | **0.83** | **0.83** | **0.82** | **0.83** | **0.82** | **0.80** | **0.82** | **0.82** | **0.86** | **0.82** | 0.79 | 0.76 |
| $C_z$ | **0.90** | **0.90** | **0.91** | **0.91** | **0.87** | **0.87** | **0.85** | **0.85** | **0.93** | **0.92** | **0.89** | **0.89** | **0.92** | **0.88** | **0.87** | **0.89** |
| Maximum resolution | | | | | | | | | | | | | | | | |
| $A$ | 0.64 | 0.64 | 0.21 | 0.16 | 0.76 | 0.76 | 0.63 | 0.63 | 0.68 | 0.67 | 0.48 | 0.48 | 0.76 | 0.73 | 0.52 | 0.47 |
| $C_x$ | **0.94** | **0.94** | **0.93** | **0.93** | **0.92** | **0.92** | **0.92** | **0.92** | **0.97** | **0.97** | **0.94** | **0.94** | **0.96** | **0.96** | **0.93** | **0.93** |
| $C_y$ | **0.86** | **0.86** | **0.75** | **0.86** | **0.82** | **0.82** | **0.80** | **0.81** | **0.80** | **0.80** | **0.82** | **0.82** | **0.84** | **0.80** | **0.77** | 0.73 |
| $C_z$ | **0.89** | **0.89** | **0.89** | **0.90** | **0.85** | **0.85** | **0.83** | **0.83** | **0.91** | **0.91** | **0.87** | **0.87** | **0.90** | **0.86** | **0.82** | **0.84** |

* Excellent: .8 ≤ ICC (dark green, bold); good: .65 ≤ ICC < .8 (light green); moderate: .5 ≤ ICC < .65 (yellow); poor: ICC<.5 (light red).

**Table S2c**. ICC values of area sizes $A$ and centroids $C = (C_x, C_y, C_z)^T$ estimated for intensity of 105% RMT of EDC using the cortical meshes with maximum resolution when ignoring non-MEP points (M1) or removing them (M2).*

| | FDI | | ADM | | APB | | FPB | | EDC | | FDS | | ECR | | FCR | |
|---|---|---|---|---|---|---|---|---|---|---|---|---|---|---|---|---|
| | M1 | M2 | M1 | M2 | M1 | M2 | M1 | M2 | M1 | M2 | M1 | M2 | M1 | M2 | M1 | M2 |
| Resolution: 15,000 vertices | | | | | | | | | | | | | | | | |
| $A$ | **0.85** | 0.79 | **0.81** | **0.74** | 0.66 | 0.71 | **0.86** | **0.87** | **0.72** | 0.70 | 0.71 | 0.69 | **0.92** | **0.90** | 0.67 | 0.69 |
| $C_x$ | **0.96** | **0.96** | **0.94** | **0.94** | **0.96** | **0.96** | **0.96** | **0.96** | **0.94** | **0.94** | **0.94** | **0.95** | **0.94** | **0.95** | **0.93** | **0.93** |
| $C_y$ | 0.69 | 0.72 | 0.62 | 0.61 | 0.77 | 0.78 | **0.85** | **0.86** | 0.78 | 0.78 | 0.66 | 0.67 | 0.77 | 0.78 | 0.76 | 0.76 |
| $C_z$ | **0.86** | **0.86** | **0.78** | 0.79 | **0.86** | **0.87** | **0.83** | **0.93** | **0.88** | **0.88** | **0.86** | **0.86** | **0.87** | **0.88** | **0.84** | **0.85** |
| Resolution: 100,000 vertices | | | | | | | | | | | | | | | | |
| $A$ | **0.87** | **0.85** | **0.65** | 0.50 | 0.67 | 0.68 | **0.89** | **0.90** | **0.39** | 0.34 | 0.52 | 0.52 | **0.85** | **0.84** | 0.38 | 0.37 |
| $C_x$ | **0.97** | **0.97** | **0.93** | **0.93** | **0.98** | **0.98** | **0.97** | **0.98** | **0.95** | **0.95** | **0.95** | **0.95** | **0.96** | **0.96** | **0.93** | **0.93** |
| $C_y$ | 0.74 | 0.74 | 0.69 | 0.75 | 0.68 | 0.67 | **0.87** | **0.87** | **0.52** | 0.77 | 0.68 | 0.66 | **0.81** | **0.81** | **0.80** | **0.80** |
| $C_z$ | **0.87** | **0.87** | **0.75** | 0.76 | **0.81** | **0.81** | **0.80** | **0.90** | **0.80** | **0.85** | **0.86** | **0.86** | **0.83** | **0.83** | **0.83** | **0.84** |
| Maximum resolution | | | | | | | | | | | | | | | | |
| $A$ | **0.86** | **0.85** | **0.66** | 0.56 | 0.72 | 0.73 | **0.86** | **0.85** | **0.37** | 0.33 | 0.53 | 0.52 | **0.87** | **0.87** | **0.37** | 0.36 |
| $C_x$ | **0.96** | **0.96** | **0.91** | **0.93** | **0.97** | **0.97** | **0.97** | **0.97** | **0.94** | **0.95** | **0.95** | **0.95** | **0.95** | **0.95** | **0.93** | **0.93** |
| $C_y$ | 0.74 | 0.74 | 0.67 | 0.71 | 0.70 | 0.71 | **0.84** | **0.84** | **0.57** | **0.80** | **0.64** | 0.63 | **0.81** | **0.79** | **0.81** | **0.81** |
| $C_z$ | **0.87** | **0.87** | **0.73** | 0.73 | **0.81** | **0.81** | **0.80** | **0.90** | **0.80** | **0.85** | **0.88** | **0.89** | **0.82** | **0.81** | **0.85** | **0.85** |

* Excellent: .8 ≤ ICC (dark green, bold); good: .65 ≤ ICC < .8 (light green); moderate: .5 ≤ ICC < .65 (yellow); poor: ICC<.5 (light red).

**Table S3a**. Outcome of the two-way ANOVA for the area sizes $A$ (in $mm^2 \cdot \mu V \cdot 10^5$) with factors of *intensity* and *session* when considering the 15,000 mesh resolution and when removing the non-MEP points (M2).*

| | $\langle A \rangle$ at 105% RMT | | | *intensity* | | *session* | | *intensity × session* | | p-value pairwise comparison | | |
|---|---|---|---|---|---|---|---|---|---|---|---|---|
| | FDI | EDC | FCR | *F* | *p* | *F* | *p* | *F* | *p* | FDI/EDC | FDI/FCR | EDC/FCR |
| FDI | 1.51±0.20 | 2.32±0.46 | 3.88±0.99 | F(2,36)=4.883 | **.03** | F(1,18)=1.128 | .30 | F(2,36)=0.163 | .71 | .16 | .06 | .27 |
| ADM | 0.72±0.13 | 0.91±0.13 | 1.46±0.35 | F(2,30)=4.921 | **.03** | F(1,15)=0.854 | .37 | F(2,30)=0.018 | .98 | .43 | .08 | .19 |
| APB | 1.55±0.41 | 1.69±0.36 | 2.96±0.89 | F(2,34)=3.412 | .07 | F(1,17)=1.819 | .20 | F(2,34)=1.903 | .18 | 1.00 | .16 | .29 |
| FPB | 1.16±0.24 | 1.43±0.31 | 1.82±0.37 | F(2,34)=3.866 | **.03** | F(1,17)=5.534 | **.03** | F(2,34)=0.627 | .48 | .35 | .09 | .46 |
| EDC | 1.00±0.14 | 1.01±0.14 | 1.44±0.26 | F(2,34)=4.115 | **.04** | F(1,17)=0.265 | .61 | F(2,34)=0.045 | .96 | 1.00 | .14 | .13 |
| FDS | 0.86±0.16 | 1.00±0.14 | 1.39±0.19 | F(2,30)=5.594 | **.01** | F(1,15)=0.142 | .71 | F(2,30)=1.163 | .32 | 1.00 | **.03** | .06 |
| ECR | 1.24±0.20 | 1.64±0.42 | 2.02±0.39 | F(2,32)=3.520 | **.04** | F(1,16)=0.716 | .41 | F(2,32)=0.241 | .79 | .48 | **.04** | .79 |
| FCR | 0.68±0.11 | 1.00±0.18 | 1.35±0.19 | F(2,32)=6.245 | **.01** | F(1,16)=0.662 | .43 | F(2,32)=1.003 | .38 | .36 | **.01** | .27 |

* Bold face implies p<.05.

**Table S3b**. Outcome of the two-way ANOVA for the area sizes $A$ (in $mm^2 \cdot \mu V \cdot 10^5$) with factors of *intensity* and *session* when considering the 100,000 mesh resolution and when removing the non-MEP points (M2).*

| | $\langle A \rangle$ at 105% RMT | | | *intensity* | | *session* | | *intensity × session* | | p-value pairwise comparison | | |
|---|---|---|---|---|---|---|---|---|---|---|---|---|
| | FDI | EDC | FCR | *F* | *p* | *F* | *p* | *F* | *p* | FDI/EDC | FDI/FCR | EDC/FCR |
| FDI | 1.48±0.19 | 2.30±0.53 | 4.69±1.52 | F(2,36)=3.959 | .06 | F(1,18)=0.84 | .37 | F(2,36)=0.372 | .56 | .25 | .12 | .31 |
| ADM | 0.71±0.15 | 0.85±0.14 | 1.74±0.52 | F(2,30)=4.261 | **.05** | F(1,15)=1.054 | .32 | F(2,30)=0.203 | .82 | 1.00 | .09 | .25 |
| APB | 1.44±0.35 | 1.75±0.41 | 3.63±1.26 | F(2,34)=3.345 | .08 | F(1,17)=1.397 | .25 | F(2,34)=1.444 | .25 | 1.00 | .17 | .34 |
| FPB | 1.11±0.24 | 1.41±0.35 | 2.30±0.67 | F(2,34)=3.035 | .09 | F(1,17)=4.891 | **.04** | F(2,34)=1.345 | .27 | .39 | .18 | .49 |
| EDC | 0.92±0.17 | 0.87±0.11 | 1.61±0.34 | F(2,34)=5.798 | **.02** | F(1,17)=0.04 | .85 | F(2,34)=0.016 | .98 | 1.00 | **.04** | .09 |
| FDS | 0.80±0.13 | 0.93±0.16 | 1.50±0.23 | F(2,30)=6.808 | **.00** | F(1,15)=1.021 | .33 | F(2,30)=1.488 | .24 | 1.00 | **.02** | .07 |
| ECR | 1.16±0.18 | 1.50±0.39 | 2.14±0.46 | F(2,30)=3.444 | **.05** | F(1,15)=0.249 | .63 | F(2,30)=0.796 | .46 | 0.88 | **.05** | .52 |
| FCR | 0.66±0.11 | 0.92±0.16 | 1.35±0.19 | F(2,30)=6.093 | **.01** | F(1,15)=1.666 | .22 | F(2,30)=1.77 | .19 | .41 | **.01** | .25 |

* Bold face implies p<.05.

**Table S4**. Outcome of the two-way ANOVA for the $C_x$ (in mm) with factors of *intensity* and *session* when considering the 15,000, 100,000 and maximum mesh resolution and when removing the non-MEP points (M2).*

| | $\langle A \rangle$ at 105% RMT | | | *intensity* | | *session* | | *intensity × session* | | p-value pairwise comparison | | |
|---|---|---|---|---|---|---|---|---|---|---|---|---|
| | FDI | EDC | FCR | *F* | *p* | *F* | *p* | *F* | *p* | FDI/ EDC | FDI/ FCR | EDC/ FCR |
| **S4a:** Resolution: 15,000 vertices | | | | | | | | | | | | |
| FDI | 21.26±2.06 | 20.71±2.03 | 20.95±1.95 | F(2,36)=0.676 | .52 | F(1,18)=0.316 | .58 | F(2,36)=1.273 | .29 | .52 | 1.00 | 1.00 |
| ADM | 21.06±2.08 | 20.81±2.06 | 20.99±2.13 | F(2,30)=0.112 | .89 | F(1,15)=0.072 | .79 | F(2,30)=4.687 | **.02** | 1.00 | 1.00 | 1.00 |
| APB | 21.02±2.06 | 20.56±2.00 | 20.42±2.09 | F(2,34)=0.901 | .42 | F(1,17)=0.008 | .93 | F(2,34)=1.468 | .25 | .78 | .65 | 1.00 |
| FPB | 20.66±1.93 | 20.56±2.04 | 20.44±1.97 | F(2,34)=0.105 | .90 | F(1,17)=0.022 | .88 | F(2,34)=0.017 | .94 | 1.00 | 1.00 | 1.00 |
| EDC | 19.74±2.00 | 19.92±2.01 | 19.87±1.91 | F(2,34)=0.115 | .89 | F(1,17)=0.688 | .42 | F(2,34)=1.001 | .38 | 1.00 | 1.00 | 1.00 |
| FDS | 21.40±2.09 | 21.07±2.22 | 21.51±2.06 | F(2,30)=0.293 | .75 | F(1,15)=0.127 | .73 | F(2,30)=0.011 | .99 | 1.00 | 1.00 | 1.00 |
| ECR | 19.86±2.06 | 19.43±2.12 | 19.40±2.14 | F(2,32)=0.631 | .54 | F(1,16)=0.229 | .64 | F(2,32)=1.054 | .36 | 1.00 | .91 | 1.00 |
| FCR | 20.32±2.15 | 20.46±1.99 | 20.82±2.12 | F(2,32)=0.365 | .70 | F(1,16)=0.093 | .76 | F(2,32)=0.871 | .43 | 1.00 | 1.00 | 1.00 |
| **S4b:** Resolution: 100,000 vertices | | | | | | | | | | | | |
| FDI | 21.62±2.09 | 20.91±1.95 | 21.03±2.03 | F(2,36)=1.618 | .21 | F(1,18)=0.044 | .84 | F(2,36)=1.734 | .19 | .16 | .71 | 1.00 |
| ADM | 21.16±2.16 | 20.95±2.06 | 21.14±2.04 | F(2,30)=0.152 | .86 | F(1,15)=0.041 | .84 | F(2,30)=1.968 | .16 | 1.00 | 1.00 | 1.00 |
| APB | 21.42±2.13 | 20.96±2.07 | 20.23±2.09 | F(2,34)=3.04 | .06 | F(1,17)=0.019 | .89 | F(2,34)=2.077 | .14 | .93 | .13 | .43 |
| FPB | 20.88±2.03 | 20.53±2.07 | 20.31±1.95 | F(2,34)=0.733 | .49 | F(1,17)=0.014 | .91 | F(2,34)=0.221 | .80 | 1.00 | .83 | 1.00 |
| EDC | 20.14±2.08 | 19.94±2.06 | 19.84±1.99 | F(2,34)=0.658 | .52 | F(1,17)=0.41 | .53 | F(2,34)=0.054 | .95 | 1.00 | .93 | 1.00 |
| FDS | 21.72±2.10 | 21.24±2.25 | 21.30±2.04 | F(2,30)=0.396 | .68 | F(1,15)=0.014 | .91 | F(2,30)=0.043 | .96 | 1.00 | 1.00 | 1.00 |
| ECR | 20.46±2.27 | 20.23±2.25 | 20.22±2.2 | F(2,30)=0.227 | .80 | F(1,15)=0.08 | .78 | F(2,30)=0.629 | .54 | 1.00 | 1.00 | 1.00 |
| FCR | 20.15±2.29 | 20.56±2.12 | 20.57±2.28 | F(2,30)=0.359 | .70 | F(1,15)=0.037 | .85 | F(2,30)=0.161 | .85 | 1.00 | 1.00 | 1.00 |
| **S4c:** Maximum resolution | | | | | | | | | | | | |
| FDI | 21.08±2.07 | 20.59±1.94 | 20.95±2.03 | F(2,36)=0.677 | .52 | F(1,18)=0.012 | .91 | F(2,36)=0.825 | .45 | .46 | 1.00 | 1.00 |
| ADM | 20.51±2.60 | 20.07±2.49 | 20.18±2.45 | F(2,24)=0.477 | .63 | F(1,12)=0.558 | .47 | F(2,24)=1.128 | .34 | 1.00 | 1.00 | 1.00 |
| APB | 21.15±2.04 | 20.58±2.02 | 20.22±2.08 | F(2,34)=1.668 | .20 | F(1,17)=0.026 | .87 | F(2,34)=1.235 | .30 | .63 | .38 | 1.00 |
| FPB | 20.59±2.07 | 20.33±2.06 | 20.28±1.97 | F(2,34)=0.238 | .79 | F(1,17)=0.002 | .97 | F(2,34)=0.153 | .86 | 1.00 | 1.00 | 1.00 |
| EDC | 19.94±2.07 | 19.50±2.03 | 19.69±1.99 | F(2,34)=0.674 | .52 | F(1,17)=0.733 | .40 | F(2,34)=0.395 | .68 | .82 | 1.00 | 1.00 |
| FDS | 21.78±2.13 | 20.94±2.20 | 21.41±2.08 | F(2,30)=1.031 | .37 | F(1,15)=0.022 | .88 | F(2,30)=0.204 | .82 | .52 | 1.00 | 1.00 |
| ECR | 20.44±2.27 | 19.81±2.20 | 20.12±2.23 | F(2,30)=0.884 | .42 | F(1,15)=0.065 | .80 | F(2,30)=0.27 | .77 | .55 | 1.00 | 1.00 |
| FCR | 20.17±2.24 | 20.22±2.11 | 20.60±2.24 | F(2,30)=0.321 | .73 | F(1,15)=0.032 | .86 | F(2,30)=0.28 | .76 | 1.00 | 1.00 | 1.00 |

* Bold face implies p<.05.

**Table S5**. Outcome of the two-way ANOVA for the $C_y$ (in mm) with factors of *intensity* and *session* when considering the 15,000, 100,000 and maximum mesh resolution and when removing the non-MEP points (M2).*

| | $\langle A \rangle$ at 105% RMT | | | *intensity* | | *session* | | *intensity × session* | | p-value pairwise comparison | | |
|---|---|---|---|---|---|---|---|---|---|---|---|---|
| | FDI | EDC | FCR | *F* | *p* | *F* | *p* | *F* | *p* | FDI/EDC | FDI/FCR | EDC/FCR |
| **S5a:** Resolution: 15,000 vertices | | | | | | | | | | | | |
| FDI | 30.94±1.27 | 29.80±1.27 | 30.23±1.05 | F(2,36)=1.026 | .37 | F(1,18)=0.176 | .68 | F(2,36)=3.644 | **.04** | .60 | 1.00 | 1.00 |
| ADM | 30.17±1.65 | 29.20±1.56 | 29.84±1.51 | F(2,30)=0.479 | .62 | F(1,15)=0.65 | .43 | F(2,30)=1.901 | .17 | 1.00 | 1.00 | 1.00 |
| APB | 31.58±1.62 | 30.00±1.41 | 30.23±1.35 | F(2,34)=2.001 | .15 | F(1,17)=0.006 | .94 | F(2,34)=3.793 | **.05** | .38 | .30 | 1.00 |
| FPB | 30.56±1.67 | 30.20±1.52 | 30.34±1.27 | F(2,34)=0.057 | .90 | F(1,17)=0.252 | .62 | F(2,34)=1.7 | .21 | 1.00 | 1.00 | 1.00 |
| EDC | 29.44±1.43 | 29.79±1.45 | 29.51±1.27 | F(2,34)=0.100 | .91 | F(1,17)=0.258 | .62 | F(2,34)=1.26 | .30 | 1.00 | 1.00 | 1.00 |
| FDS | 29.77±1.41 | 28.73±1.51 | 28.97±1.20 | F(2,30)=0.658 | .53 | F(1,15)=0.041 | .84 | F(2,30)=0.57 | .57 | 1.00 | 1.00 | 1.00 |
| ECR | 29.55±1.61 | 29.19±1.64 | 29.11±1.36 | F(2,32)=0.144 | .87 | F(1,16)=0.255 | .62 | F(2,32)=2.316 | .12 | 1.00 | 1.00 | 1.00 |
| FCR | 29.68±1.48 | 29.38±1.37 | 28.63±1.25 | F(2,32)=0.587 | .56 | F(1,16)=0.058 | .81 | F(2,32)=0.764 | .47 | 1.00 | .68 | 1.00 |
| **S5b:** Resolution: 100,000 vertices | | | | | | | | | | | | |
| FDI | 31.32±1.41 | 30.07±1.36 | 30.87±1.09 | F(2,36)=1.226 | .31 | F(1,18)=0.04 | .84 | F(2,36)=3.657 | **.04** | .45 | 1.00 | .99 |
| ADM | 30.27±1.86 | 29.55±1.64 | 30.35±1.53 | F(2,30)=0.426 | .66 | F(1,15)=0.196 | .66 | F(2,30)=1.713 | .20 | 1.00 | 1.00 | 1.00 |
| APB | 31.78±1.66 | 30.94±1.46 | 30.56±1.41 | F(2,34)=0.905 | .41 | F(1,17)=0.068 | .80 | F(2,34)=2.137 | .13 | 1.00 | .58 | 1.00 |
| FPB | 30.64±1.73 | 30.33±1.49 | 30.46±1.35 | F(2,34)=0.046 | .90 | F(1,17)=0.069 | .80 | F(2,34)=1.071 | .33 | 1.00 | 1.00 | 1.00 |
| EDC | 29.73±1.42 | 30.30±1.48 | 29.58±1.28 | F(2,34)=0.487 | .62 | F(1,17)=0.652 | .43 | F(2,34)=2.032 | .15 | 1.00 | 1.00 | .97 |
| FDS | 30.34±1.44 | 29.49±1.55 | 29.64±1.15 | F(2,30)=0.538 | .59 | F(1,15)=0.104 | .75 | F(2,30)=0.812 | .45 | .96 | 1.00 | 1.00 |
| ECR | 28.83±1.61 | 29.12±1.68 | 29.06±1.31 | F(2,30)=0.058 | .94 | F(1,15)=0.109 | .75 | F(2,30)=2.469 | .10 | 1.00 | 1.00 | 1.00 |
| FCR | 29.46±1.38 | 29.84±1.52 | 29.15±1.27 | F(2,30)=0.32 | .73 | F(1,15)=0.044 | .84 | F(2,30)=0.241 | .79 | 1.00 | 1.00 | 1.00 |
| **S5c:** Maximum resolution | | | | | | | | | | | | |
| FDI | 31.01±1.41 | 29.79±1.36 | 30.28±1.12 | F(2,36)=0.994 | .38 | F(1,18)=0.773 | .39 | F(2,36)=4.401 | **.02** | .42 | 1.00 | 1.00 |
| ADM | 30.23±2.25 | 29.23±1.99 | 29.64±1.76 | F(2,24)=0.488 | .62 | F(1,12)=2.171 | .17 | F(2,24)=2.021 | .15 | 1.00 | 1.00 | 1.00 |
| APB | 31.22±1.65 | 30.58±1.49 | 30.20±1.40 | F(2,34)=0.555 | .58 | F(1,17)=0.05 | .83 | F(2,34)=2.018 | .15 | 1.00 | .96 | 1.00 |
| FPB | 30.46±1.75 | 30.44±1.56 | 30.17±1.34 | F(2,34)=0.041 | .96 | F(1,17)=0.63 | .44 | F(2,34)=0.596 | .56 | 1.00 | 1.00 | 1.00 |
| EDC | 29.50±1.47 | 30.07±1.55 | 29.29±1.24 | F(2,34)=0.478 | .62 | F(1,17)=0.799 | .38 | F(2,34)=1.721 | .19 | 1.00 | 1.00 | 1.00 |
| FDS | 29.74±1.55 | 28.93±1.62 | 28.85±1.24 | F(2,30)=0.468 | .63 | F(1,15)=0.575 | .46 | F(2,30)=1.016 | .37 | 1.00 | 1.00 | 1.00 |
| ECR | 28.63±1.63 | 28.68±1.74 | 28.87±1.33 | F(2,30)=0.042 | .96 | F(1,15)=0.211 | .65 | F(2,30)=3.445 | **.05** | 1.00 | 1.00 | 1.00 |
| FCR | 29.12±1.40 | 29.64±1.62 | 29.03±1.27 | F(2,30)=0.260 | .77 | F(1,15)=0.012 | .91 | F(2,30)=0.795 | .46 | 1.00 | 1.00 | 1.00 |

* Bold face implies p<.05.

**Table S6**. Outcome of the two-way ANOVA for the $C_z$ (in mm) with factors of *intensity* and *session* when considering the 15,000, 100,000 and maximum mesh resolution and when removing the non-MEP points (M2).*

| | $\langle A \rangle$ at 105% RMT | | | *intensity* | | *session* | | *intensity × session* | | p-value pairwise comparison | | |
|---|---|---|---|---|---|---|---|---|---|---|---|---|
| | FDI | EDC | FCR | *F* | *p* | *F* | *p* | *F* | *p* | FDI/EDC | FDI/FCR | EDC/FCR |
| **S6a:** Resolution: 15,000 vertices | | | | | | | | | | | | |
| FDI | 108.18±1.69 | 108.65±1.72 | 107.70±1.55 | F(2,36)=1.097 | .35 | F(1,18)=0.065 | .80 | F(2,36)=2.316 | .11 | 1.00 | 1.00 | .60 |
| ADM | 109.11±1.80 | 109.14±1.93 | 108.56±1.72 | F(2,30)=0.380 | .69 | F(1,15)=0.144 | .71 | F(2,30)=0.714 | .50 | 1.00 | 1.00 | 1.00 |
| APB | 108.19±1.60 | 108.75±1.58 | 108.07±1.52 | F(2,34)=0.563 | .53 | F(1,17)=0.035 | .85 | F(2,34)=1.420 | .26 | 1.00 | 1.00 | 1.00 |
| FPB | 109.12±1.70 | 108.14±1.74 | 108.09±1.71 | F(2,34)=1.031 | .34 | F(1,17)=0.317 | .58 | F(2,34)=0.849 | .44 | 1.00 | .35 | 1.00 |
| EDC | 109.40±1.66 | 108.55±1.80 | 108.78±1.65 | F(2,34)=0.929 | .37 | F(1,17)=0.244 | .63 | F(2,34)=1.099 | .35 | .89 | .18 | 1.00 |
| FDS | 110.07±1.65 | 109.42±1.89 | 109.55±1.66 | F(2,30)=0.602 | .49 | F(1,15)=0.004 | .95 | F(2,30)=0.356 | .70 | 1.00 | .41 | 1.00 |
| ECR | 108.92±1.85 | 108.81±1.70 | 108.68±1.66 | F(2,32)=0.054 | .95 | F(1,16)=0.012 | .91 | F(2,32)=0.777 | .41 | 1.00 | 1.00 | 1.00 |
| FCR | 110.35±1.46 | 109.51±1.68 | 109.87±1.57 | F(2,32)=0.741 | .45 | F(1,16)=0.164 | .69 | F(2,32)=0.732 | .49 | .96 | .85 | 1.00 |
| **S6b:** Resolution: 100,000 vertices | | | | | | | | | | | | |
| FDI | 108.70±1.67 | 109.35±1.79 | 107.99±1.56 | F(2,36)=2.154 | .15 | F(1,18)=0.074 | .79 | F(2,36)=3.139 | .06 | 1.00 | .37 | .23 |
| ADM | 109.82±1.85 | 109.73±1.95 | 108.75±1.80 | F(2,30)=1.312 | .28 | F(1,15)=0.003 | .96 | F(2,30)=1.274 | .29 | 1.00 | .55 | .56 |
| APB | 108.90±1.71 | 109.61±1.66 | 108.51±1.57 | F(2,34)=1.060 | .36 | F(1,17)=0.079 | .78 | F(2,34)=1.572 | .22 | 1.00 | 1.00 | .43 |
| FPB | 109.83±1.67 | 109.13±1.79 | 108.54±1.72 | F(2,34)=1.245 | .29 | F(1,17)=0.000 | .98 | F(2,34)=0.367 | .70 | 1.00 | .22 | 1.00 |
| EDC | 110.42±1.66 | 109.35±1.86 | 109.35±1.67 | F(2,34)=1.557 | .23 | F(1,17)=0.324 | .58 | F(2,34)=0.505 | .61 | .67 | **.04** | 1.00 |
| FDS | 110.80±1.63 | 110.44±1.86 | 109.79±1.61 | F(2,30)=1.207 | .30 | F(1,15)=0.002 | .96 | F(2,30)=0.542 | .59 | 1.00 | .06 | 1.00 |
| ECR | 110.86±1.52 | 110.37±1.73 | 110.11±1.60 | F(2,30)=0.65 | .53 | F(1,15)=0.112 | .74 | F(2,30)=0.577 | .57 | 1.00 | .32 | 1.00 |
| FCR | 110.99±1.57 | 110.19±1.78 | 110.15±1.67 | F(2,30)=0.897 | .42 | F(1,15)=0.071 | .79 | F(2,30)=0.281 | .76 | 0.88 | .32 | 1.00 |
| **S6c:** Maximum resolution | | | | | | | | | | | | |
| FDI | 108.60±1.71 | 109.26±1.80 | 108.02±1.53 | F(2,36)=1.558 | .23 | F(1,18)=0.177 | .68 | F(2,36)=2.879 | .07 | 1.00 | .87 | .39 |
| ADM | 108.92±2.15 | 108.98±2.31 | 108.54±2.13 | F(2,24)=0.262 | .77 | F(1,12)=0.366 | .56 | F(2,24)=1.155 | .33 | 1.00 | 1.00 | 1.00 |
| APB | 108.92±1.70 | 109.51±1.66 | 108.47±1.57 | F(2,34)=0.822 | .45 | F(1,17)=0.048 | .83 | F(2,34)=0.844 | .44 | 1.00 | 1.00 | .65 |
| FPB | 109.78±1.70 | 109.03±1.80 | 108.59±1.69 | F(2,34)=1.103 | .33 | F(1,17)=0.022 | .88 | F(2,34)=0.137 | .87 | 1.00 | .26 | 1.00 |
| EDC | 110.22±1.66 | 109.14±1.84 | 109.34±1.62 | F(2,34)=1.411 | .26 | F(1,17)=0.281 | .60 | F(2,34)=0.389 | .68 | .56 | .19 | 1.00 |
| FDS | 110.75±1.63 | 110.46±1.86 | 109.96±1.62 | F(2,30)=0.663 | .47 | F(1,15)=0.075 | .79 | F(2,30)=0.959 | .40 | 1.00 | .21 | 1.00 |
| ECR | 110.76±1.53 | 110.35±1.75 | 109.72±1.52 | F(2,30)=1.242 | .30 | F(1,15)=0.128 | .73 | F(2,30)=1.568 | .23 | 1.00 | .13 | 1.00 |
| FCR | 110.95±1.54 | 110.19±1.80 | 110.15±1.66 | F(2,30)=0.810 | .42 | F(1,15)=0.138 | .72 | F(2,30)=0.199 | .82 | 1.00 | .22 | 1.00 |

* Bold face implies p<.05.

**Table S7**. Outcome of the two-way ANOVA for the amplitude (μV) with factors of *intensity* and *session* when considering the 15,000 mesh resolution and when removing the non-MEP points (M2).*

| | $\langle A \rangle$ at 105% RMT | | | *intensity* | | *session* | | *intensity × session* | | p-value pairwise comparison | | |
|---|---|---|---|---|---|---|---|---|---|---|---|---|
| | FDI | EDC | FCR | *F* | *p* | *F* | *p* | *F* | *p* | FDI/ EDC | FDI/ FCR | EDC/ FCR |
| FDI | 286.59±38.83 | 365.52±64.20 | 561.15±127.18 | F(2,36)=5.913 | **.02** | F(1,18)=0.513 | .48 | F(2,36)=0.520 | .50 | .20 | .05 | .12 |
| ADM | 183.30±34.68 | 187.29±34.84 | 260.86±56.30 | F(2,30)=5.372 | **.03** | F(1,15)=0.064 | .80 | F(2,30)=0.792 | .46 | 1.00 | .07 | .12 |
| APB | 343.04±84.23 | 299.11±55.85 | 407.51±104.16 | F(2,34)=1.191 | .32 | F(1,17)=1.401 | .25 | F(2,34)=0.645 | .53 | 1.00 | 1.00 | .44 |
| FPB | 236.32±38.56 | 223.33±38.80 | 278.20±53.89 | F(2,34)=2.053 | .16 | F(1,17)=7.836 | **.01** | F(2,34)=2.315 | .11 | 1.00 | .70 | .25 |
| EDC | 170.86±20.53 | 160.85±12.49 | 210.73±21.32 | F(2,36)=6.413 | **.00** | F(1,18)=0.626 | .44 | F(2,36)=0.352 | .71 | 1.00 | .07 | **.01** |
| FDS | 168.04±19.51 | 179.02±23.05 | 222.47±27.17 | F(2,30)=4.572 | **.02** | F(1,15)=3.047 | .10 | F(2,30)=1.604 | .22 | 1.00 | .09 | .07 |
| ECR | 222.46±23.31 | 247.82±38.31 | 276.44±37.80 | F(2,32)=2.586 | .09 | F(1,16)=0.320 | .58 | F(2,32)=0.237 | .72 | .99 | .07 | .75 |
| FCR | 157.21±13.97 | 174.40±18.41 | 203.78±20.27 | F(2,32)=4.645 | **.02** | F(1,16)=0.068 | .80 | F(2,32)=0.420 | .66 | .72 | .07 | .13 |

* Bold face implies p<.05.

**Table S8**. The ICC values for measurement are shown in the table for all the muscles. Int 1,2,3 represent the intensities of 105% RMT of FDI, EDC and FCR, respectively. The bold font implies that ICC values are excellent (ICC>0.8).

| **FDI** | **Int1** | **Int2** | **Int3** | **ADM** | **Int1** | **Int2** | **Int3** | **APB** | **Int1** | **Int2** | **Int3** | **FPB** | **Int1** | **Int2** | **Int3** |
|---|---|---|---|---|---|---|---|---|---|---|---|---|---|---|---|
| Amp (μV) | **0.85** | **0.90** | 0.52 | Amp (μV) | 0.65 | 0.76 | **0.86** | Amp (μV) | 0.74 | 0.74 | **0.82** | Amp (μV) | 0.66 | **0.93** | **0.84** |
| Lat (ms) | **0.90** | **0.96** | **0.93** | Lat (ms) | **0.88** | **0.78** | **0.80** | Lat (ms) | 0.79 | 0.58 | **0.84** | Lat (ms) | 0.30 | 0.71 | 0.27 |
| CoGx (mm) | **0.96** | **0.97** | **0.97** | CoGx (mm) | **0.93** | **0.93** | **0.95** | CoGx (mm) | **0.94** | **0.96** | **0.96** | CoGx (mm) | **0.92** | **0.96** | **0.97** |
| CoGy (mm) | **0.85** | 0.78 | 0.74 | CoGy (mm) | 0.73 | 0.77 | 0.73 | CoGy (mm) | **0.86** | 0.76 | 0.79 | CoGy (mm) | **0.80** | **0.84** | 0.72 |
| CoGz (mm) | **0.95** | **0.91** | **0.90** | CoGz (mm) | **0.92** | **0.88** | **0.87** | CoGz (mm) | **0.91** | **Int1** | **Int2** | **Int3** | **0.87** | **0.93** | **0.89** |
| **EDC** | **Int1** | **Int2** | **Int3** | **FDS** | **Int1** | **Int2** | **Int3** | **ECR** | **Int1** | **Int2** | **Int3** | **FCR** | **Int1** | **Int2** | **Int3** |
| Amp (μV) | **0.87** | 0.72 | **0.87** | Amp (μV) | **0.83** | **0.85** | 0.58 | Amp (μV) | **0.91** | **0.97** | **0.86** | Amp (μV) | 0.70 | 0.75 | 0.47 |
| Lat (ms) | 0.68 | 0.57 | **0.87** | Lat (ms) | 0.64 | 0.64 | **0.81** | Lat (ms) | **0.82** | **0.90** | **0.91** | Lat (ms) | 0.63 | **0.84** | **0.86** |
| CoGx (mm) | **0.97** | **0.96** | **0.97** | CoGx (mm) | **0.96** | **0.96** | **0.95** | CoGx (mm) | **0.96** | **0.96** | **0.97** | CoGx (mm) | **0.96** | **0.94** | **0.97** |
| CoGy (mm) | **0.86** | 0.66 | 0.77 | CoGy (mm) | **0.84** | 0.77 | 0.70 | CoGy (mm) | **0.86** | **0.83** | 0.78 | CoGy (mm) | **0.81** | **0.83** | 0.79 |
| CoGz (mm) | **0.95** | **0.85** | **0.90** | CoGz (mm) | **0.92** | **0.93** | **0.87** | CoGz (mm) | **0.95** | **0.87** | **0.91** | CoGz (mm) | **0.90** | **0.90** | **0.95** |
